# Spontaneous formation of prebiotic compartment colonies on Hadean Earth and pre-Noachian Mars

**DOI:** 10.1101/2021.05.11.443509

**Authors:** Elif S. Köksal, Inga Põldsalu, Henrik Friis, Stephen Mojzsis, Martin Bizzarro, Irep Gözen

## Abstract

The primitive cells that emerged at the origin of life are commonly viewed as spherical biosurfactant shells, freely suspended in aqueous media^1–3^. This model explains initial, but not subsequent events in the development process towards structured protocells. Taking into consideration the involvement of naturally occurring surfaces, which were abundant on the early Earth^4^, we report feasible and productive pathways for the development of primitive cells. Surfaces intrinsically possess energy, easily utilized by the interfacing amphiphiles, such as lipids, to attain self-organization and spontaneous transformations^5–7^. We show that the physical interaction of phospholipid pools with 20 Hadean Earth analogue materials as well as a Martian meteorite composed of fused regolith representing the ancient crust of Mars, consistently lead to the shape transformation and autonomous formation of surfactant compartment assemblies. Dense, colony-like protocell populations grow from these lipid deposits, predominantly at the grain boundaries or cleavages of the investigated natural surfaces, and remain there for several days. The model protocells in our study are able to autonomously develop, transform and pseudo-divide, and encapsulate RNA as well as DNA. We also demonstrate that they can accommodate non-enzymatic, DNA strand displacement reactions. Our findings suggest a feasible route towards the transformation from nonliving to living entities, and provide fresh support for the ‘Lipid World’ hypothesis^8^.

---

All contemporary life forms feature cell membranes, two-dimensional fluid films composed of biosurfactants, which define the boundaries of the cell, and those of its internal substructures. Evidence suggests that the protocell building blocks, chemically closely related to today’s membrane constituents, may have existed prior to 4 billion years ago in sufficient concentrations to allow for the assembly of extended membranous structures; either via synthesis from simpler precursor molecules under prebiotic conditions^9–12^, or by meteorite delivery via late accretion^13,14^. Yet, how featureless self-assembled containers -considered to be the stepping stone to life-may have developed on the early Earth, and what their exact structural and dynamic characteristics were, remains undefined. The current paradigm calls for the involvement of spherical primitive soft compartments which formed by self-assembly in bulk aqueous environments^1–3^. Biosurfactant assemblies are, however, capable of amazingly versatile shape and phase transformations, requiring only tiny energy input. Our recent work shows that through the involvement of an artificial solid surface, strikingly sophisticated sequences of autonomous soft matter shape transformations are possible^15–17^. A potentially critical role of natural surfaces in enabling the formation and development of protocells has possibly been overlooked all along, until interactions of amphiphiles with solid mineral microparticles under formation of closed compartments was reported^18–21^. The possible presence of lipid molecules and the abundance of rocks and minerals on the early Earth and Mars, provides a strong incentive for transferring the previously observed spontaneous vesiculation on engineered surfaces to natural specimens. This represents a critical step in the understanding of protocell formation and fate, which would significantly extent the ‘lipid world’ hypothesis.

Here, we focus now on the impact of the natural surfaces on formation and development of model protocells. We utilize 20 thin, continuous flat natural Earth surfaces, i.e. rocks, minerals, glasses (*cf.* **Table S1** for a complete list), and a Martian meteorite specimen, and exposed them to lipid reservoirs. The Martian meteorite sample, Northwest Africa 7533 (NWA 7533), is documented to contain ancient crustal fragments that date back to the earliest history of the planet. The centimeter-sized optically transparent micro-thin sections provide extended areas for direct observation of topological transformations. The polished surfaces are not uniformly flat; they retain some of the irregularities (cracks, fissures, grain boundaries) of the natural materials.

We report results from a total of 21 naturally occurring surfaces (**Table S1**) prepared as substrates for lipid membrane adhesion. For microscopy imaging, the surfaces were planarized to an even thickness of 170 μm (**Fig. 1a**). **Fig. 1b** shows, a top view photograph of a semitransparent granite section, and a largely opaque section of the Martian meteorite NWA 7533.

**Figure 1.**
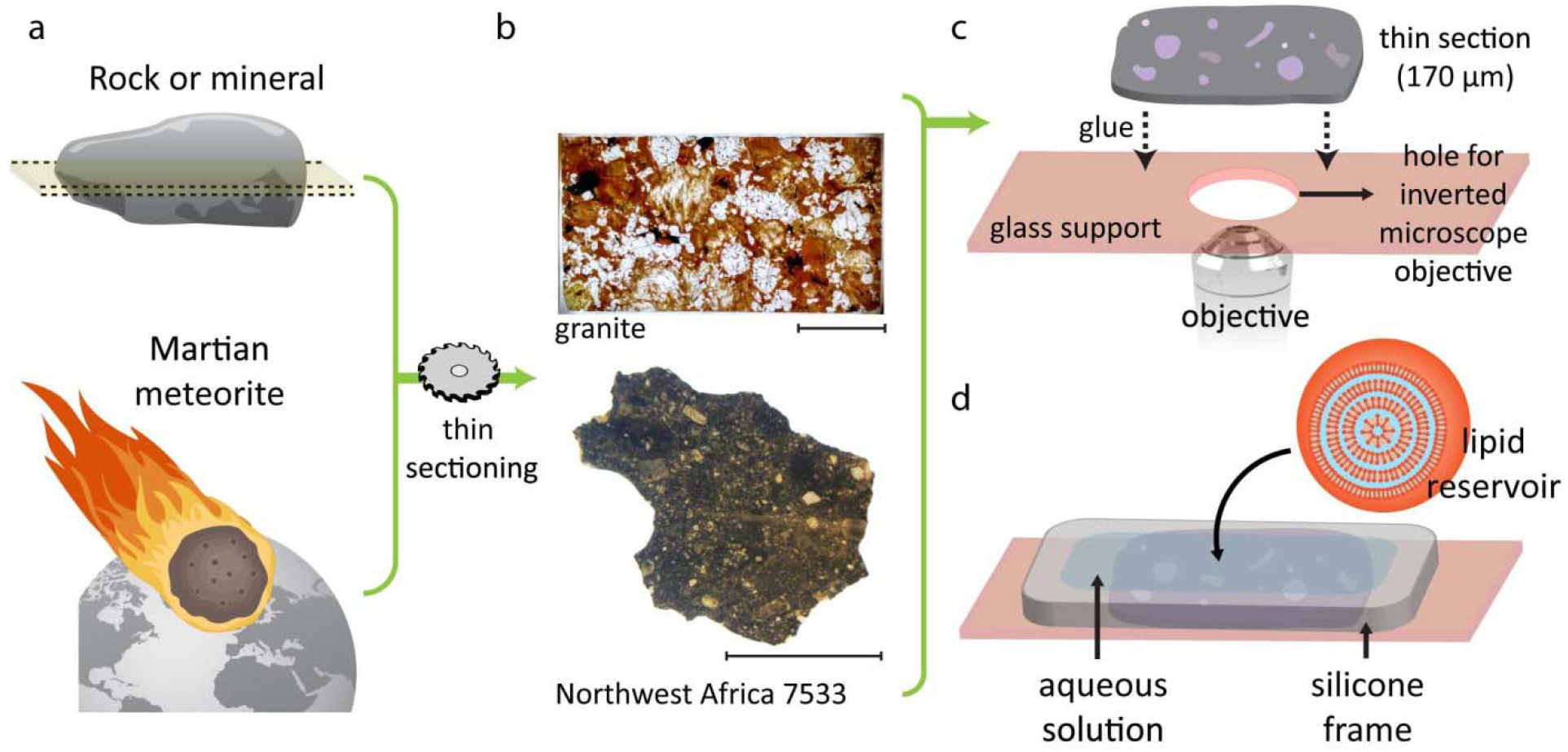
Summary of the experiment. **a**, Earth surfaces (20 specimen) and one Martian meteorite NWA 7533 sample have been planarized to an even thickness of 170 μm. **b**, photographs of two examples of natural surfaces: granite (top) and NWA 7533 (bottom), from top view. Scale bars: 1 cm. (***cf.* Fig. S1** for photographs of other specimens) **c**, the schematic drawing showing the preparation of the sample chamber. The thin section has been gently mounted on a perforated glass cover slide (ID 0.5 cm). The hole allows direct contact with the oil objective of an inverted confocal microscope. The glass slide provides support for the thin fragile sections. **d**, A flat-bottomed silicon frame is air-sealed onto the glass cover slip and filled with an aqueous suspension of lipid reservoirs. For almost completely opaque samples, a standard cover slide with an upright confocal microscope was employed (not shown).

For inverted confocal fluorescence microscopy of fully- or semi-transparent surfaces, the thin sections were mounted on a perforated glass cover slip (**Fig. 1c**). Opaque samples were mounted on a standard cover slip, and an upright fluorescence confocal microscope was used instead. A silicone frame was adhered to the glass carrier, surrounding the specimen, and filled with aqueous solution. Multilamellar lipid reservoirs in aqueous suspension were added to the solution with an automatic pipette.

Upon contact of the lipid reservoir with the surface, wetting and subsequent transformations occur, resulting in the formation of the adherent lipid compartments physically connected via a dense lipid nanotube network (tube ∅~100 nm). We have recently thoroughly characterized and reported these transformations on nano-engineered silica substrates^15,16^, and reproduced these experiments for this study (**Fig. 2a**). Briefly, each reservoir spontaneously spreads as a double lipid bilayer membrane, followed by rupture of the distal (upper) bilayer, and its rapid transformation into a network of nanotubes. Sections of the nanotubes swell over time to form vesicular compartments, governed by the tendency of the system to reduce its bending energy cost, minimizing the surface free energy of the membrane. **Fig. 2a** shows a lipid nanotube network with small vesicles nucleating from the network on a deposited SiO_2_ surface. **Fig. 2b** is a schematic drawing of a single nanotube-connected vesicle as seen in panel (**a**). **Fig. 2c-f** shows confocal micrographs of vesicle-nanotube networks on several natural mineral, glass and rock surfaces: olivine, fluorite, dolerite and picrite basalt. SI contains additional examples (**Fig. S2, Section 11**), including sequences of earlier steps of the transformation described above, e.g. rupturing of double bilayers (**Fig. S3 SI**). Protocell populations formed via the nanotube-mediated path are typically physically connected to each other, but can also separate from the population, migrate to remote locations, and adhere there^15^. These features of the nanotube-vesicle networks led to a new early division hypothesis^22^.

**Figure 2.**
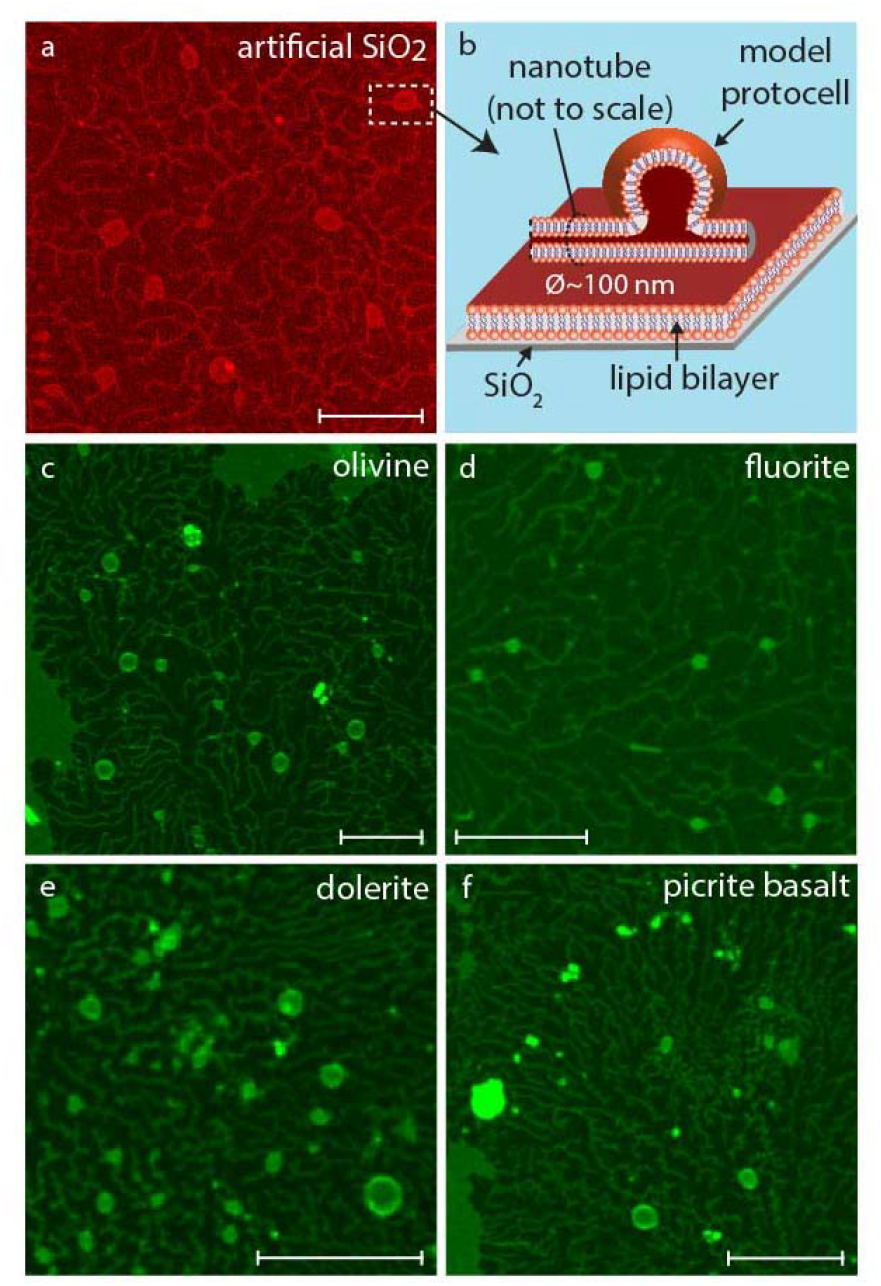
Formation of lipid nanotube-protocell networks. **a**, confocal micrograph showing the positive control: a lipid nanotube-protocell network formed on a nano-engineered SiO_2_ surface, from top view. **b**, schematic drawing depicting the membrane configuration of the structure shown in a white dashed frame. A nanotube, a fraction of which is swelling as a lipid compartment, resides on a single bilayer. **c-f**, Confocal micrographs showing the phenomenon described in (a-b) on natural surfaces: **c**, olivine. **d**, fluorite. **e**, dolerite. **f**, picrite basalt. Scale bars: 10 μm.

Besides the heterogeneous composition of natural surfaces with distinct grain boundaries, a prominent difference to synthetic surfaces is the presence of fissures and cracks. We observe the emergence of protocell colonies predominantly along these natural interface features (**Fig. 3,** also **Fig. S4**). The figure shows examples of protocell colonies emerging on olivine (**a**), oligoclase (**b**), eclogite (**c**), nephelinite (**d**), granite (**e**), eclogite (**f**) and fluorite (**e**). Panels **(a)**, **(d)** and **(g)** are epi-fluorescence projections of the colonies while **(b)**, **(e)** and **(i)** are cross sections (x-y plane). **(c)** and **(f)** show cross sections along the dashed lines in **(b)**, **(e)** and **(h)** (x-z plane). In **Fig. 3c,** up to three stacked layers of protocells are distinguishable. **Fig. 3f** (also **Fig. S5, S11**) reveals the presence of subcompartments inside single, dome-shaped protocells, a phenomenon we had recently reported on artificial surfaces^17^. In the report, we pointed out a possible pathway to pseudo-division as the consequence of rupture of the enveloping surfactant shell.

**Figure 3.**
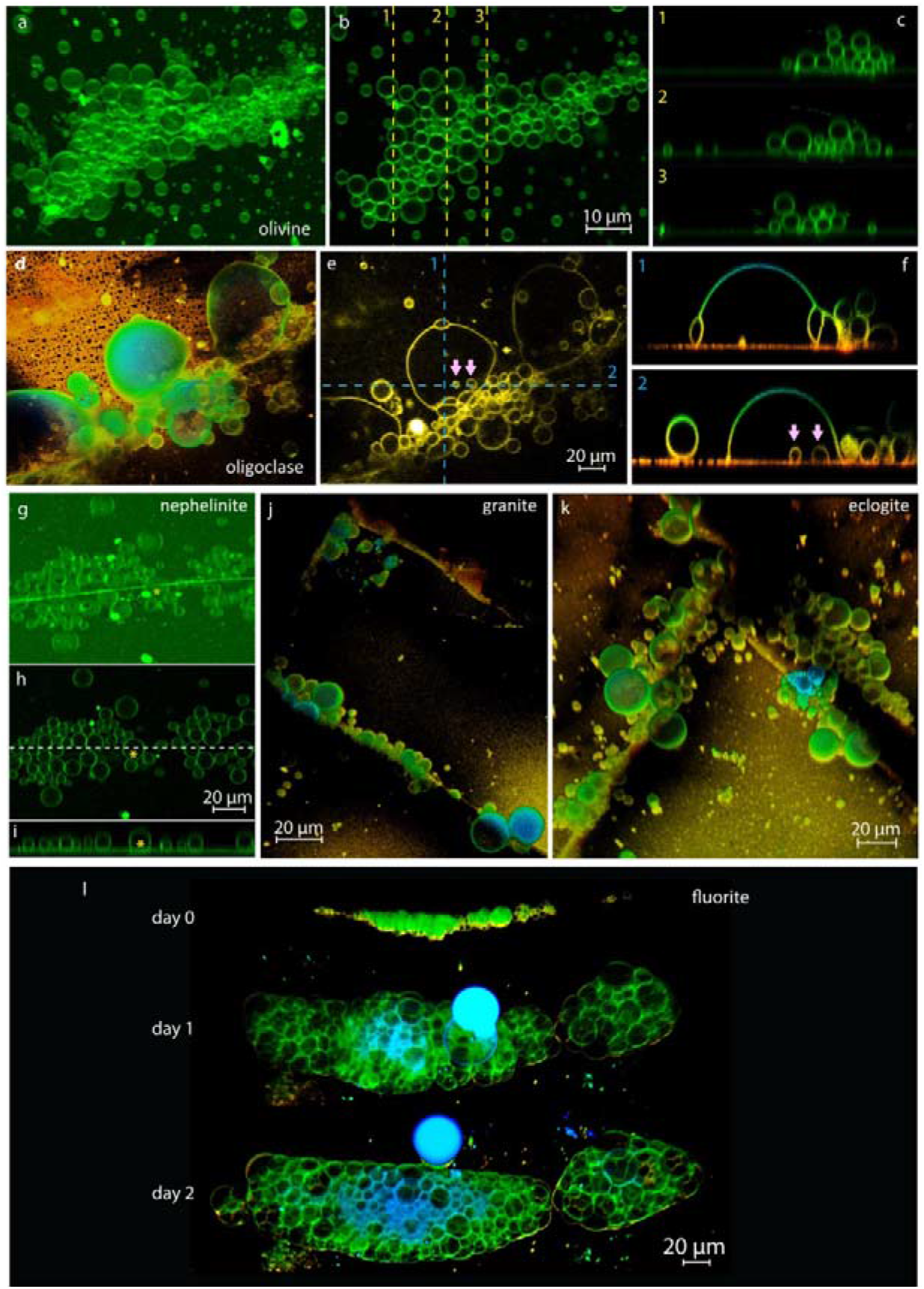
Formation of protocell colonies on analogue Hadean materials. Confocal micrographs showing dense, protocell colonies, formed predominantly along the cracks and grain boundaries of the surfaces: **a-c,** olivine. **d-f,** oligoclase. **g-i,** nephelinite. Panels (a), (d) and (g) are epi-fluorescence projections of the colonies while (b), (e) and (i) are cross sections (x-y plane). (c), (f) and (i) show cross sections along the dashed lines in (b), (e) and (h), respectively (x-z plane). **j,** granite. **k,** eclogite. **l,** fluorite. The foam-like colonies grows in 2 days. Arrows in (e) and (f) point to encapsulated containers (surface-adhered “sub-compartments”).

In minerals the cleavages and fractures follow specific orientations dictated by the crystal structure. This results in a change in surface energy and wettability depending on the surface exposed, as each surface orientation will have a slightly different atomic density^23,24^. The cleavages and fractures represent areas where the atomic density is changed resulting in a local change of surface energy. For the rock samples, which contain aggregates of multiple minerals with different orientations, each grain in the rock will have its own surface energy, but this is abruptly changed at the grain boundary. Neighboring minerals in the rocks will all have different orientations, even between grains of the same mineral. Consequently, every grain boundary presents a slightly different surface energy regime. SEM-EDX scans of several of the Earth materials showing the element composition maps of the surfaces are provided in **SI Section S7**. The analyses reveal that the regions amplifying the protocell formation on these substrates occur predominantly and consistently (but not exclusively) at the interface of quartz-containing grains.

Volcanic and impact generated glasses are thought to have been present on the pre-Noachian Mars^25^ and on the Hadean Earth^26^. The glass samples we used were freshly synthesized standard analogues to natural surfaces matching the chemical compositions of terrestrial crustal rocks. X-ray fluorescence (XRF) spectroscopy-based compositional analyses of the rock glasses are provided in **SI Section 8**.

The possibility of the existence of vesicle colonies on the early Earth surfaces has been hypothesized in earlier work^27^. Colonies increase the mechanical stability and provide an advantage for segregating, fusing and sharing internalized compounds. The close proximity of colony vesicles can likely facilitate direct chemical exchange, e.g. through transient pores^17^.

The same experimental procedure was performed on the Martian meteorite NWA 7533, which contains fragments of the planet’s earliest crust^28,29^. Recent investigation of the elemental and isotopic compositions of these fragments indicate that the early crust of Mars was reworked by impacts in the presence of water, establishing that water was present on Mars >4.4 billion years ago^30^. Water, minerals and organic molecules may have led to early life on Mars, similar to its emergence on the early Earth. Evidence for extraterrestrial existence of suitable phospholipids and fatty acids exists, as these molecules have been extracted from meteorites earlier^13,14^.

The meteorite sample is significantly more fragile and rougher, compared to all other surfaces used in this study. On this sample, we also observe the formation of foam-like vesicle colonies (**Fig. 4a**). Two cross sectional views (x-z plane) over the numbered lines indicated with white dashed lines in **(a)**, have been inserted into the top right corner of the micrograph with corresponding numbering. In **Fig. 4b** another region with several colonies is depicted. Over four days, the foam-like structures develop into smaller colonies, each with multiple protocells (*cf.* SI for another example).

**Figure 4.**
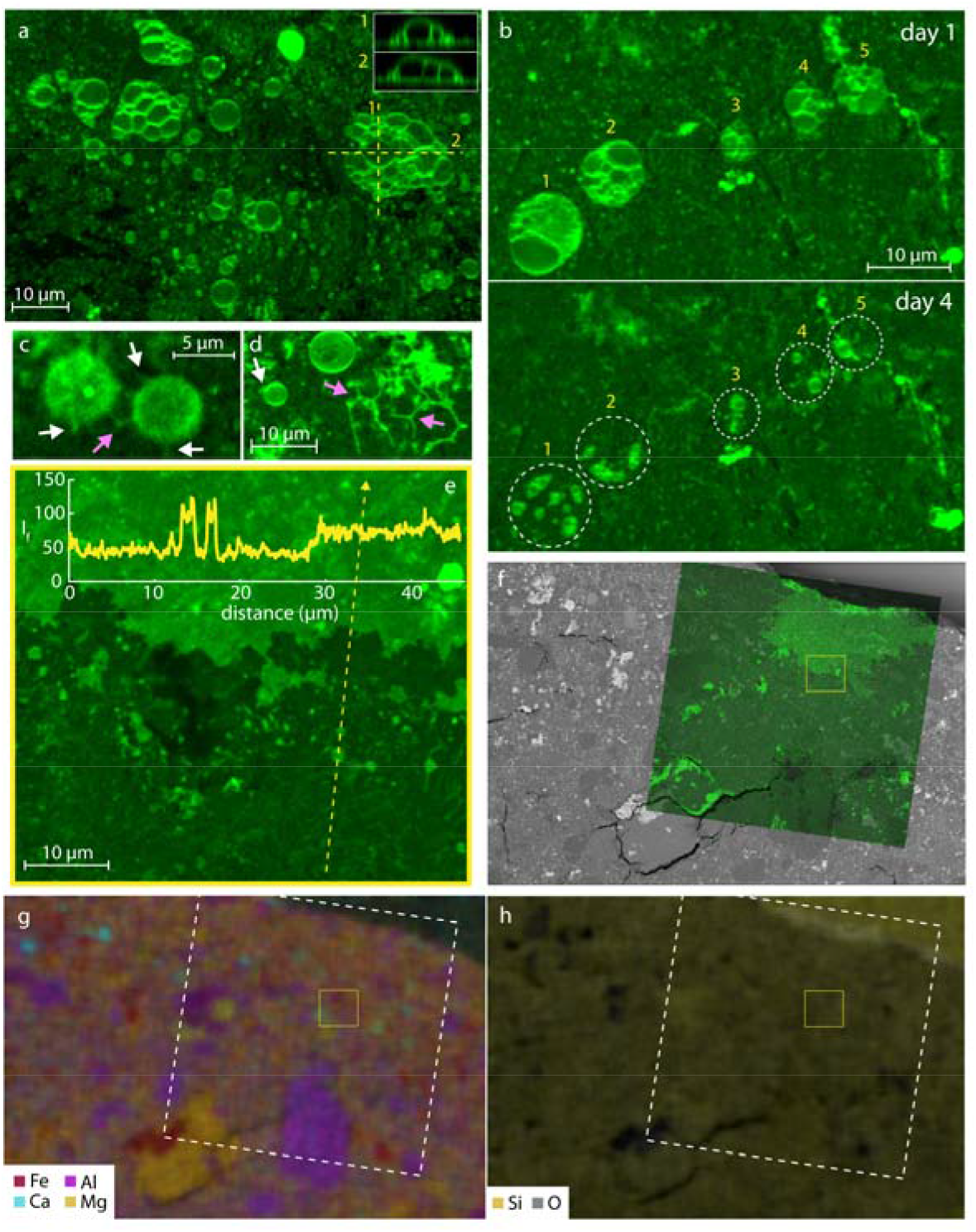
Formation of protocells on the Martian meteorite NWA 7533 specimen. **a**, confocal micrograph showing several foam-like protocell colonies emerging on the proximal (lower) bilayer that is adhered onto the meteorite. The insets show the cross sectional profile of the colonies (xz plane) along the dashed lines. **b**, Confocal micrographs showing the division of the colonies after 4 days to smaller units. The original structures and their corresponding daughter structures have been labeled with identical numbers. **c-d**, close up confocal sections showing the nanotubular structures associated with the vesicles (white arrows). Pink arrows show the Y-junctions, 3-way tubular intersections typically associated with lipid nanotube networks. **e**, confocal micrograph showing a double lipid bilayer (bright green) dewetting a single lipid bilayer (dark green), and small, nanotube-attached vesicles adhered on the bilayer. The inset plot shows the fluorescence intensity profile over the dashed line. The two M-shaped spikes correspond to the two vesicles on the single bilayer. **f**, Superimposed fluorescence micrograph of the membrane and SEM image of the region it is positioned on the meteorite. Yellow frame represents (e). **g-h**, SEM-EDX scans of the meteorite surface shown in (f) showing different elements. White dashed lines correspond to the contour of the fluorescence membrane micrograph in (f).

Vesicles stemming from the nanotubes are depicted in **Fig. 4c-d** (white arrows). They show the typical structure of freely suspended lipid nanotubes, featuring intersections of three conduits with including 120 degree angles (Y-junctions). The Y-junctions are the energetically most favorable configuration of branching lipid nanotubes^31^.

**Fig. 4e** shows a membrane region with a ruptured distal bilayer edge, separating single (darker green) and double (brighter green) bilayer. An intensity profile along the yellow dashed line is overlaid onto the micrograph. The two M-shaped spikes are typical fluorescence intensity line profiles for fluorescently labeled vesicles. The vesicles are residing on the proximal single bilayer, distinguishable by its fluorescence intensity of approximately half the intensity of a double bilayer. A video showing the rupturing process on the meteorite specimen is available as online material. We performed scanning electron microscopy with energy-dispersive X-ray spectroscopy (SEM-EDX) of the surface to map the composition to elucidate possible influences on the protocell formation (**Fig 4f-h**). In **Fig. 4f** the fluorescence micrograph of the lipid membrane is superimposed on the SEM image. The yellow frame represents the region shown in panel (**e**). In (**g**) the distinct domains of Fe (red color), Al (magenta), Ca (cyan) and Mg (yellow) are emphasized. A particular preference for a specific element is not discernible. (**h**) shows the distribution of Si and O which are, except for a few dense regions, abundant throughout the surface. The lipid membrane spreads over the entire area on the surface of the meteorite. Additional micrographs showing vesicle formation, SEM–EDX maps of several regions on the meteorite surface and point analyses of distinct domains are supplied in **SI Section 9**.

To evaluate the uptake of genetic components by the model protocells described above, we superfused vesicles and vesicle colonies adhered to the natural surfaces, e.g. granite, with RNA solution and observed spontaneous encapsulation (**Fig. 5a-e**). Briefly, a region of interest of ~150 μm in diameter on the sample surface was selected, and vesicles in that region were superfused with fluorescently labeled RNA over a period of up to 4 minutes. The fluorescence signal in the internal volume of one compartment (magenta circle in panel **d**) as well as in a region outside but nearby the vesicle (gray circle in panel **d**) were monitored with laser scanning confocal fluorescence microscopy. Emission intensities of membrane (red channel) and labeled RNA (green channel) were recorded before (**Fig. 5b**), during (**Fig. 5c**) and after (**Fig. 5d**) exposure to the RNA, as shown in panel (**e**). White dashed lines outline the containers in the RNA channel, red and white solid circles are the regions of interest (ROI) of intra- and extravesicular space. The concentration inside the vesicle reaches 80% of the concentration of the stock solution of the RNA in the ambient solution within ~ 2 minutes of superfusion. After this exposure, the protocellular compartment slowly leaks its fluorescent contents, but maintains 50% of the initial ambient concentration for several minutes (*cf.* **Fig. S21** for another example). The transmembrane uptake occurs directly via transient membrane pores, which we characterized in earlier studies^16,17^.

**Figure 5.**
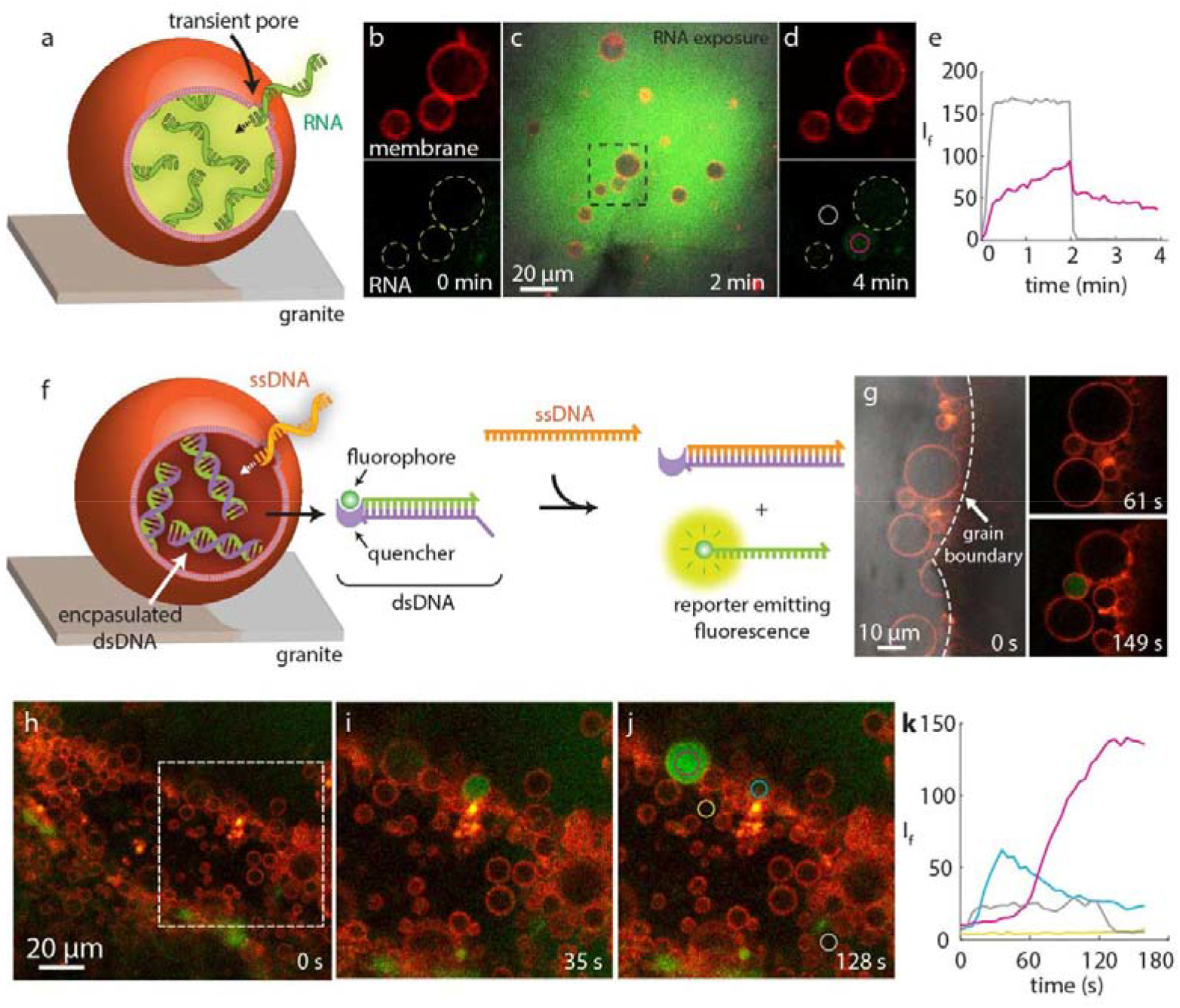
RNA and DNA encapsulation and non-enzymatic DNA strand displacement reaction inside the protocells formed on natural surfaces. **a**, schematic drawing showing the RNA encapsulation via transient pores. **b-d**, confocal micrographs showing protocells (red fluorescence) before (b), during (c) and after (d) RNA (green fluorescence) superfusion. (b) and (d) represent the area framed in black dashed lines in (c). **e**, fluorescence intensity of the regions of interest in (d) (circles with continuous lines) over time. The center vesicle takes up the RNA and maintains it over 4 minutes. The sharp drop in ambient RNA intensity corresponds to the termination of controlled superfusion. **f**, schematic drawing showing the encapsulation of ssDNA as part of a non-enzymatic, entropy-driven DNA reaction. The entry of the ssDNA leads to melting of annealed dsDNA previously encapsulated inside the protocells and hybridization of one of the strands with the newly entering ssDNA (strand displacement). The released ssDNA becomes no longer quenched and starts to fluoresce. **g-j**, confocal micrographs corresponding to (f). **(i-j)** represents the region framed in white dashed line in (h). **k**, fluorescence intensity inside the ROIs circled in j, versus time. The vesicles and colonies shown in this figure were formed on granite.

We also studied the ability of the protocells formed on natural surfaces to host a non-enzymatic, DNA strand displacement reaction, which was earlier reported by Winfree et al.^32^ **(Fig. 5f**). Briefly, annealed double stranded DNA (dsDNA) fragments were added to the ambient solution of maturing vesicle colonies and incubated with them for 24 hours for encapsulation (**Fig. 5f**). One of the strands of dsDNA features a fluorophore (green color), the complementary strand its quencher (purple color), preventing emission. The incubated vesicles are then superfused with single-stranded DNA (ssDNA), complementary to the quencher strand (orange color). The ssDNA enters some of the protocells in a fashion to similar RNA (**Fig. 5a-e**), resulting in protein-free, entropy driven strand displacement and generation of a fluorescence signal from the released ssDNA. **Fig. 5g** shows an experiment in which the internal volume of a small vesicle becomes visibly fluorescent within a few minutes of ssDNA exposure of a group of preincubated vesicles. **Fig. 5h-k** shows this process on an extended surface region. **Fig. 5i-j** show a magnified region of the protocell colony in **Fig. 5h** (framed in white dashed lines). The fluorescence intensity in 4 different regions of interest was monitored during ssDNA exposure (circles in **Fig. 5j**): The reaction occurs inside at least two of the protocells in the colony. In one instance, the reaction proceeds rapidly but the product appears to leak immediately (ROI and plot in blue). In the other instance, the fluorescence signal increases (ROI and plot in magenta) over 3 minutes (*cf***. Fig. S22** for 4 additional examples).

We show that the surfaces of various naturally occurring materials promote the consistent formation of lipid compartments, which are able to encapsulate small molecules and host a DNA strand displacement reaction. Biosurfactant film structures formed on mineral, glass and rock slices utilize the surface energy for a double bilayer wetting/de-wetting process with subsequent spontaneous shape transformations, leading to the formation of closed containers. In some instances, subcompartments spontaneously emerge inside such containers at the solid interface.

We hereby provide evidence supporting a hypothesis that surfaces might have been of fundamental importance not only for the formation of robust protocells, but also for their initial development stages on the Hadean Earth. Moreover, we observe colony formation, which has been the subject of a prior hypothesis arguing for increased stability and facilitated chemical exchange in comparison to isolated single containers. While vesicle colonies were observed in earlier experimental work utilizing poly-lysine^33^, we observed spontaneous colony formation without the need for promoters. The preferential population of cracks and fissures in the specimen indicate that rough surfaces do not prevent but enhance compartment formation. We conclude that the early Earth bulk hypothesis should be expanded to include solid natural surfaces as possible key enabling factor, and suggest to not only further explore the transformation capabilities of the surface energy-driven protocell model, but to also aim to combine it with more sophisticated prebiotic chemistry.

