## Supplementary information for "Spontaneous formation of prebiotic compartment colonies on Hadean Earth and pre-Noachian Mars"

**This PDF file includes:**

Supplementary text

S1. List of all utilized surfaces and summary of associated observations
S2. Extended Figure 1: Photographs of sectioned rocks and minerals
S3. Rupture modes and bilayer analysis
S4. Extended Figure 2: Lipid nanotube-vesicle networks
S5. Extended Figure 3: Protocell colonies
S6. Subcompartmentalization
S7. SEM-EDX analysis
S7.1. Olivine
S7.2. Fluorite
S7.3. Olivine-rich lava rock
S7.4. Eclogite
S7.5. Granite
S7.6. Martian meteorite NWA7533
S8. X-ray fluorescence spectroscopy and ICP-MA analysis
S9. Extended Figure 4: Protocell formation on the NWA7533
S10. Extended Figure 5: RNA and DNA encapsulation
S11. Heat-induced protocell formation
S12. Supplementary movies

Tables S1

Figures S1 to S23

Captions for movies S1 to S2

References for SI

**Other supplementary materials for this manuscript include the following:**

Movies S1 to S2

### S1. List of utilized surfaces and summary of associated observations

**Table S1: Complete list of utilized surfaces.** Formation of: double bilayer (DB), single bilayer (SB), nanotubes (N), vesicles (V), dome-shaped vesicular compartments (D), vesicle growth on nanotubes (Nt), vesicle growth on surface fractures/cracks (Cr), floral ruptures (Fl), fractal ruptures (Fr), isolated vesicle without any connection to nanotubes or colonies (IV) (not shown).

| # | Material | Locality | Ruptures | Formations | Vesicle localization |
|---|---|---|---|---|---|
| 1 | Quartz $SiO_2$ | Hardangervidda, Norway | Fl, Fr | DB, SB, N, V, D | Nt, Cr |
| 2 | Olivine $(Mg,Fe)_2SiO_4$ | Åheim, Norway | Fl, Fr | DB, SB, N, V | Nt, Cr |
| 3 | Oligoclase $NaAlSi_3O_8$ | Bamble, Norway | Fl | DB, SB, N, V, D | Nt, Cr, IV |
| 4 | Granite | Drammen, Norway | Fl, Fr | DB, SB, N, V, D | Cr, IV |
| 5 | Eclogite | Western Gneiss Region, Norway | Fl, Fr | DB, SB, N, V, D | Nt, Cr |
| 6 | Fluorite $CaF_2$ | Østfold pegmatite field, Norway | Fl, Fr | DB, SB, N, V, D | Nt, Cr, IV |
| 7 | Calcite $CaCO_3$ | Kjørholt Mine, Norway | Fl | DB, SB, D | - |
| 8 | Olivine-rich lava rock | Hawaii, USA | Fl, Fr | DB, SB, D, N, V | Nt, Cr |
| 9 | Muscovite $KAl_2[AlSi_3O_{10}](OH)_2$ | Norway | Fl, Fr | DB, SB | IV |
| 10 | Marble | Unknown | Fl, Fr | DB, SB | IV |
| 11 | Basalt (BRP-1G) | n/a | Fl, Fr | DB, SB, N, V | Nt |
| 12 | Dolerite (DNC-1G) | n/a | Fl, Fr | DB, SB, N, V, D | Nt |
| 13 | Nephelinite (NKT-1G) | n/a | Fl, Fr | DB, SB, N, V, D | Cr |

| | | | | | |
|---|---|---|---|---|---|
| 14 | Eifel picro basalt (BKWE-1g) | n/a | Fl, Fr | DB, SB, N, V, D | Nt, Cr |
| 15 | Picrite basalt (BOOS-1G) | n/a | Fl, Fr | DB, SB, N, V | Nt, IV |
| 16 | Norite (BSWR-1G) | n/a | Fl, Fr | DB, SB, N, V | Nt |
| 17 | Diabase (W-2G) | n/a | Fl, Fr | DB, SB, N, V | Nt, IV |
| 18 | Picrite basalt (BCR-2Ga) | n/a | Fl, Fr | DB, SB, N, V, D | IV, IV |
| 19 | Andesite (AGV-2G) | n/a | Fl, Fr | DB, SB, N, V, D | Nt, IV |
| 20 | Gabbro (GSM-1G) | n/a | Fl, Fr | DB, SB, N, V | Nt, IV |
| 21 | Meteorite (NWA 7533) | Sahara Desert, Africa | Fl, Fr | DB, SB, V | Nt, Cr, IV |

**S2. Extended Figure 1: Photographs of sectioned rocks and minerals**

Natural specimens were planarized to an even thickness of 170 µm for microscopy imaging as described in **Fig.1**. **Fig. S1** shows photographs of sectioned Earth rocks and minerals from top view. Transparent and semi-transparent surfaces allow visualization with an inverted confocal microscope while opaque samples are visualized using an upright confocal microscope.

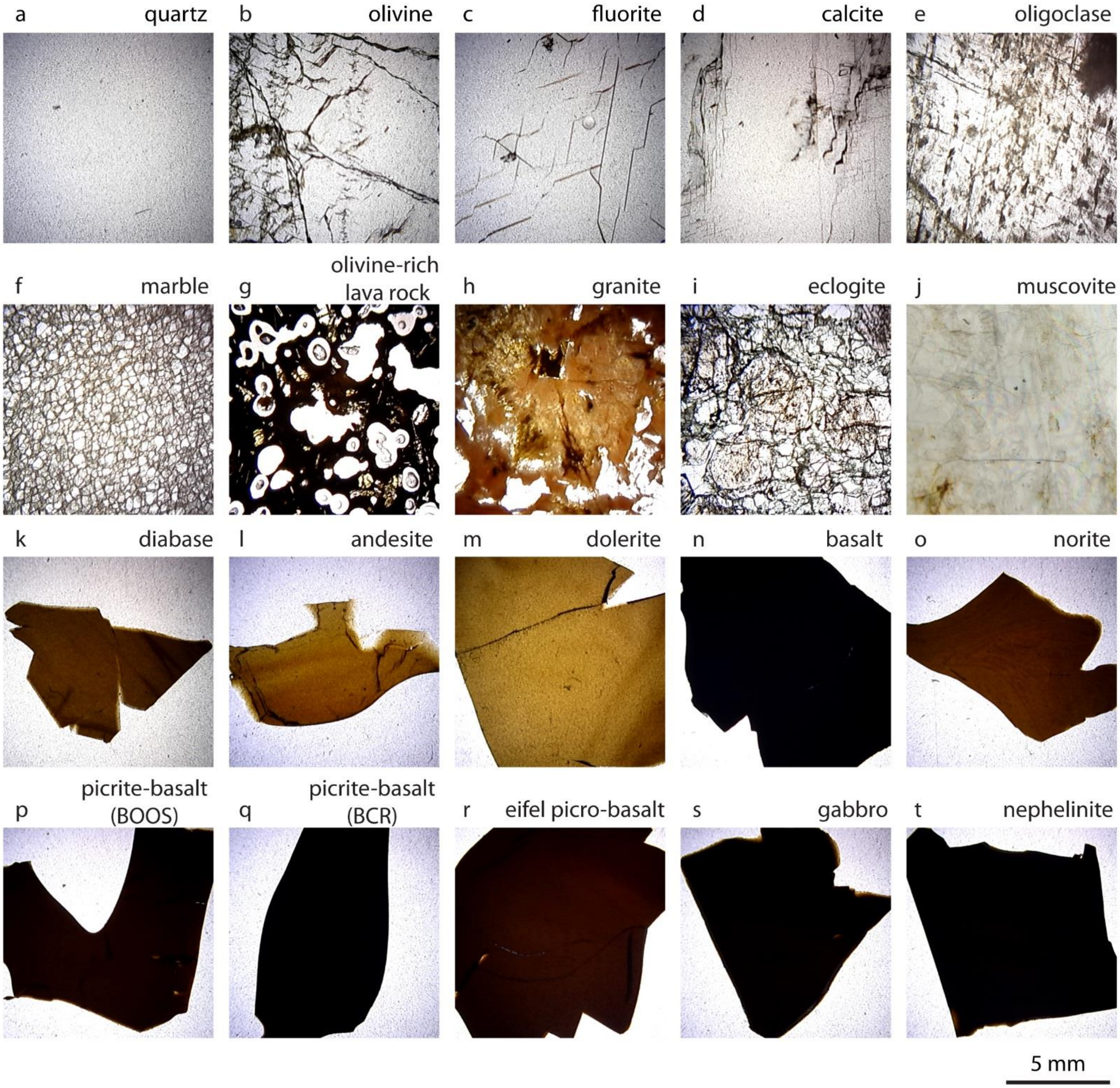


**Figure S1. Photographs of all sectioned surfaces from top view. (a)** quartz, **(b)** olivine, **(c)** fluorite, **(d)** calcite, **(e)** oligoclase, **(f)** marble, **(g)** olivine-rich lava rock, **(h)** granite, **(i)** eclogite, **(j)** muscovite**, (k)** diabase, **(l)** andesite, **(m)** dolerite, **(n)** basalt, **(o)** norite, **(p)** picrite basalt (BOOS), **(q)** picrite basalt (BCR), **(r)** eifel picro-basalt, **(s)** gabbro, **(t)** nephelinite.

### S3. Extended Figure 2: Lipid nanotube-vesicle networks

More examples of lipid nanotube vesicle networks on natural surfaces (**Fig. 2)** are shown in **Fig. S2**.

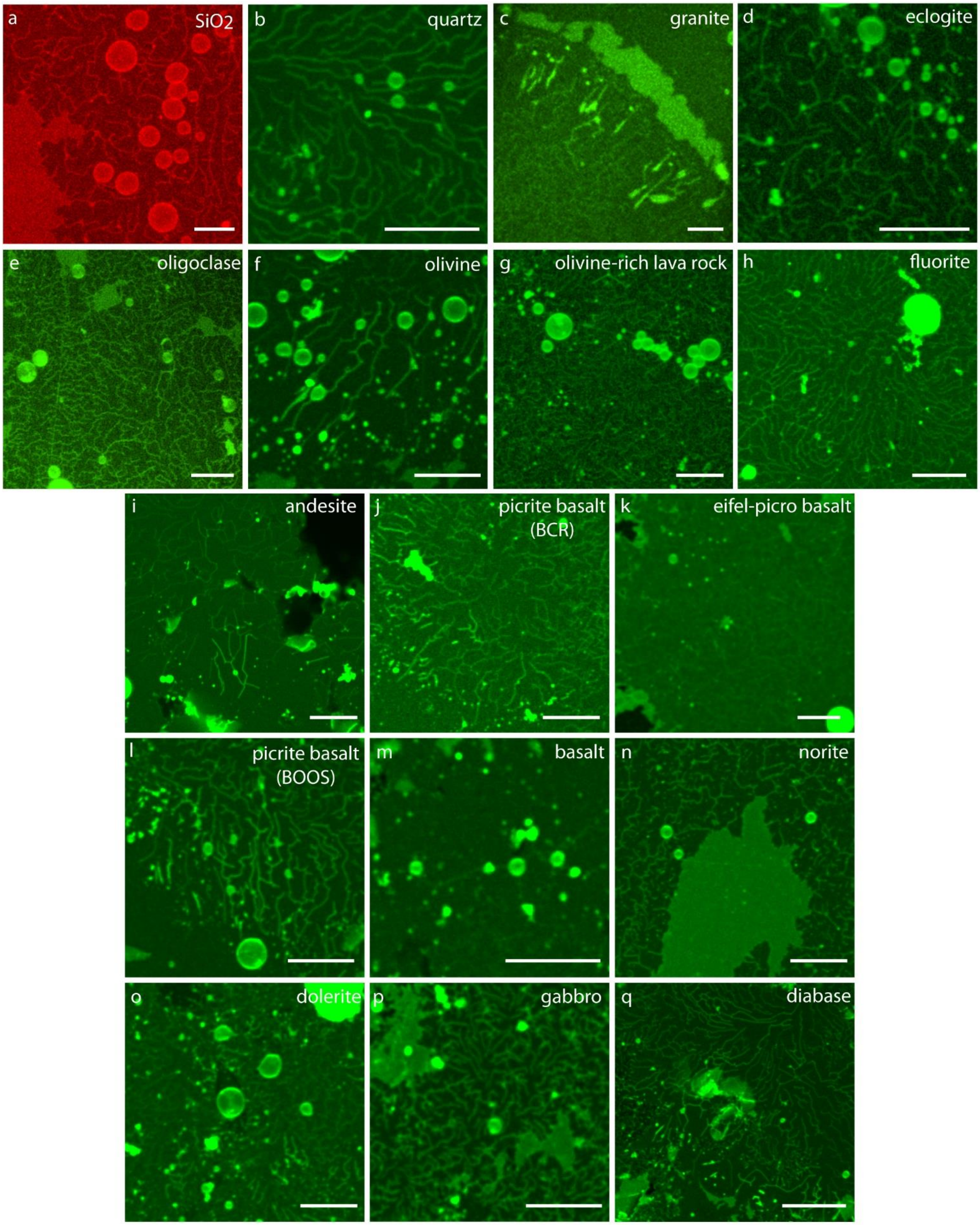


**Figure S2. Formation of lipid nanotube-vesicle networks on Hadean Earth surfaces.** Epifluorescence projection of a confocal micrograph showing a lipid nanotube-vesicle network on a single bilayer on **(a)** the positive control: a nano-engineered $SiO_2$ surface, **(b)** quartz, **(c)** granite, **(d)** eclogite, **(e)** oligoclase, **(f)** olivine, **(g)** olivine-rich lava rock, **(h)**

fluorite, **(i)** andesite, **(j)** picrite basalt (BCR), **(k)** eifel picro-basalt, **(l)** picrite basalt (BOOS), **(m)** basalt, **(n)** norite, **(o)** dolerite, **(p)** gabbro, **(q)** diabase. Scale bars: 10 µm.

### S4. Bilayer analysis and ruptures

Prior to the formation of nanotubes and emergence of protocells (**Fig.2** and **Fig. S2**), lipid reservoirs self-spread on the solid support as a double bilayer membrane. Due to continuous adhesion to the substrate, tension in the membrane increases, leading to rupturing and retraction of the distal (upper) bilayer on the proximal (lower) bilayer. Two different rupture morphologies; floral and fractal, were previously reported as a result of the transformation described above on $SiO_2$ substrates[1]. The rupture morphology is determined by the density of adhesion between the bilayers enabled by the divalent ions present in the aqueous environment, e.g. $Ca^{2+}$[2]. Double- and single- bilayers as well as fractal and floral ruptures were also observed on many natural substrates. Yellow and pink arrows in **Fig. S3** points to the fractal ruptures. The fluorescence intensity profiles along the dashed lines in **Fig. S3** reveals an approximately 2:1 intensity ratio for DB:SB.
We recently showed the rapid nucleation and growth of protocells from the lipid nanotube networks described above, induced by mild heat gradients (days vs. minutes). We have similar observations on natural surfaces which we present in **Fig. S23** in **SI Section 11**.

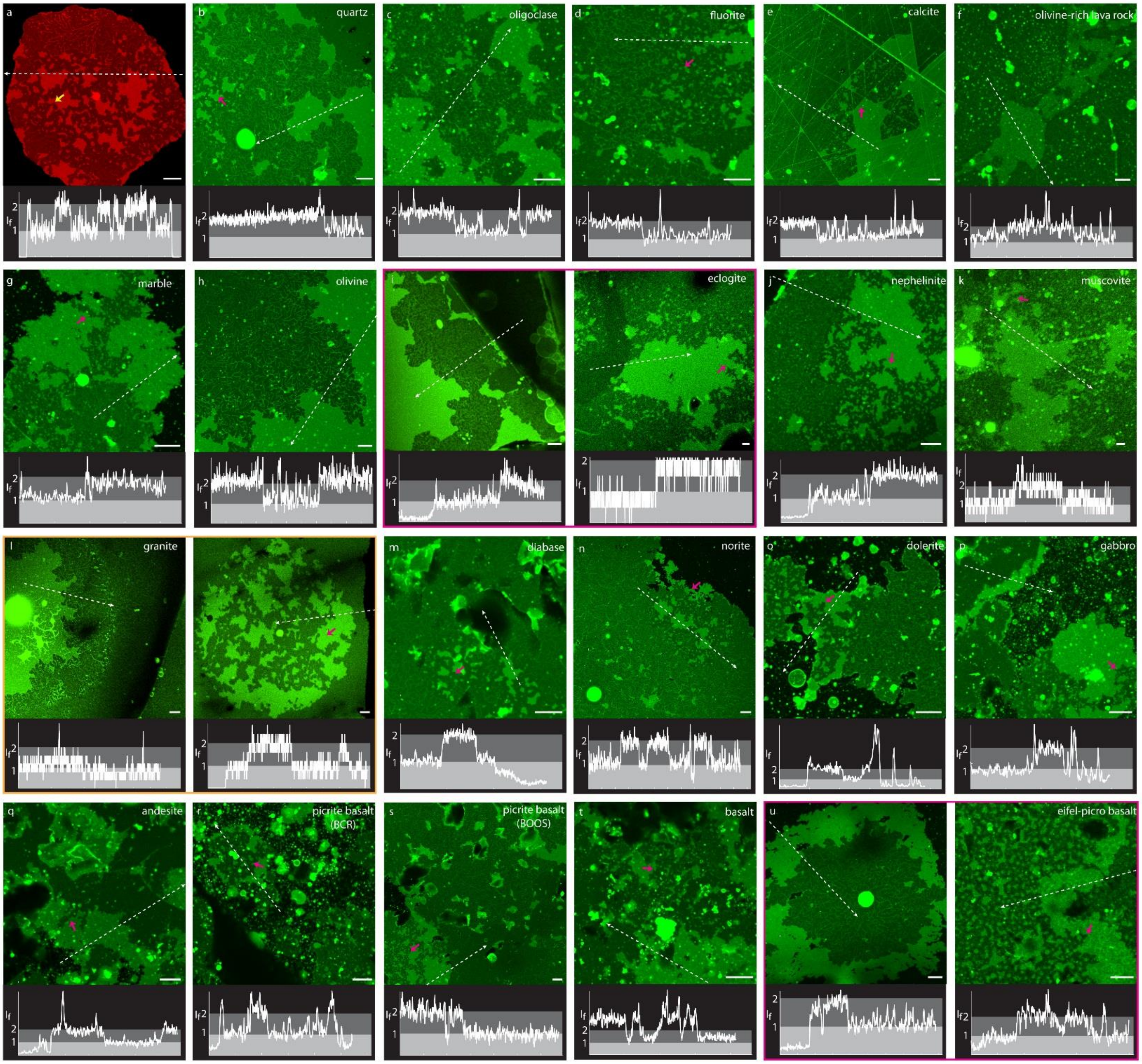


**Figure S3. Membrane lamellarity analysis and different rupture modes.** Confocal micrographs showing double- (bright green) and single-lipid bilayers (dark green). Plots below every micrograph show the fluorescence intensity profile over the dashed lines in the micrograph. The surfaces on which DLB and SLB are observed: **(a)** $SiO_2$, **(b)** quartz, **(c)** oligoclase, **(d)** fluorite, **(e)** calcite, **(f)** olivine-rich lava rock, **(g)** marble, **(h)** olivine, **(i)** eclogite, **(j)** nephelinite, **(k)** muscovite, **(l)** granite, **(m)** diabase, **(n)** norite, **(o)** dolerite, **(p)** gabbro, **(q)** andesite, **(r)** picrite basalt (BCR), **(s)** picrite basalt (BOOS), **(t)** basalt, **(u)** eifel-picro basalt. Yellow arrows in (a) and pink arrows in (b-u) show fractal type of ruptures on membranes. Scale bars: 10 µm.

### S5. Extended Figure 3: Protocell colonies

More examples of protocell colonies along the fissures/cracks shown in **Fig. 3** are provided in **Fig. S4**.

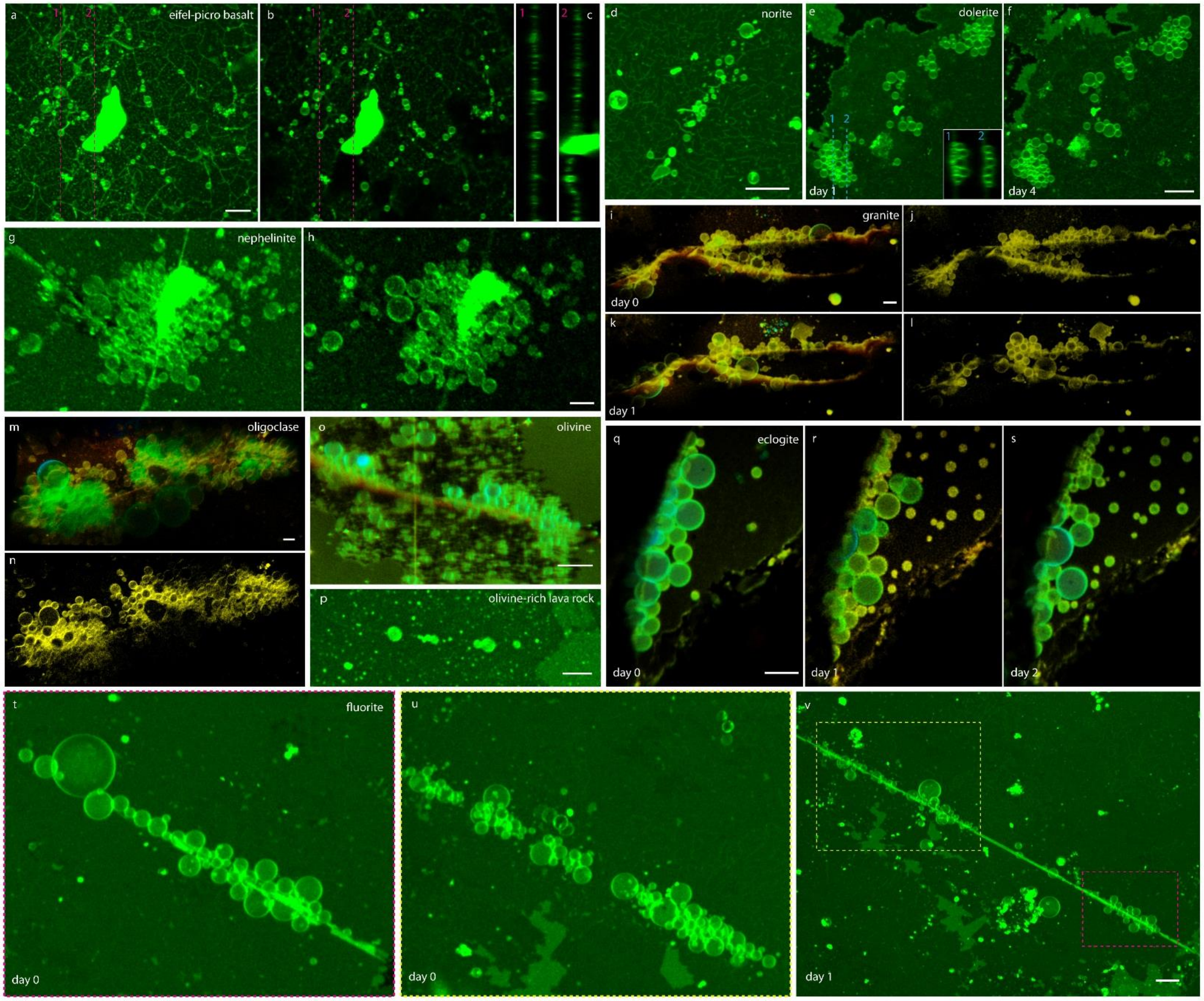


**Figure S4. Formation of protocell colonies on Earth surfaces.** Confocal micrographs showing dense protocell colonies, formed predominantly along the grain boundaries and cleavages of the surfaces: **(a-c)** eifel picro-basalt, **(d)** norite, **(e-f)** dolerite, **(g-h)** nephelinite, **(i-l)** granite, **(m-n)** oligoclase, **(o)** olivine, **(p)** olivine-rich lava rock, **(q-s)** eclogite, **(t-v)** fluorite. Panels (a), (d), (e-f), (g), (i) and (k), (m), (o), (p), (q-s) and (t-v) are epifluorescence projections of the colonies while (b), (h), (j) and (l), (n) are cross sections in (x-y plane). (c) and inset in (e) show cross sections along the dashed lines in (b) and (e), respectively (x-z plane). (e-f), (i-l), (q-s) and (t-v) show changes in the colony over days. Pink and yellow dashed frames in (v) correspond to the areas shown in (t) and (u) in day 1, respectively. Scale bars: 20 µm.

### S6. Subcompartmentalization

More examples of subcompartmentalization of protocells (**Fig. 3f)** are provided in **Fig. S5**. In case of rupturing of the enveloping protocell, subcompartments can transform to daughter protocells depicting a possible form of primitive division[3].

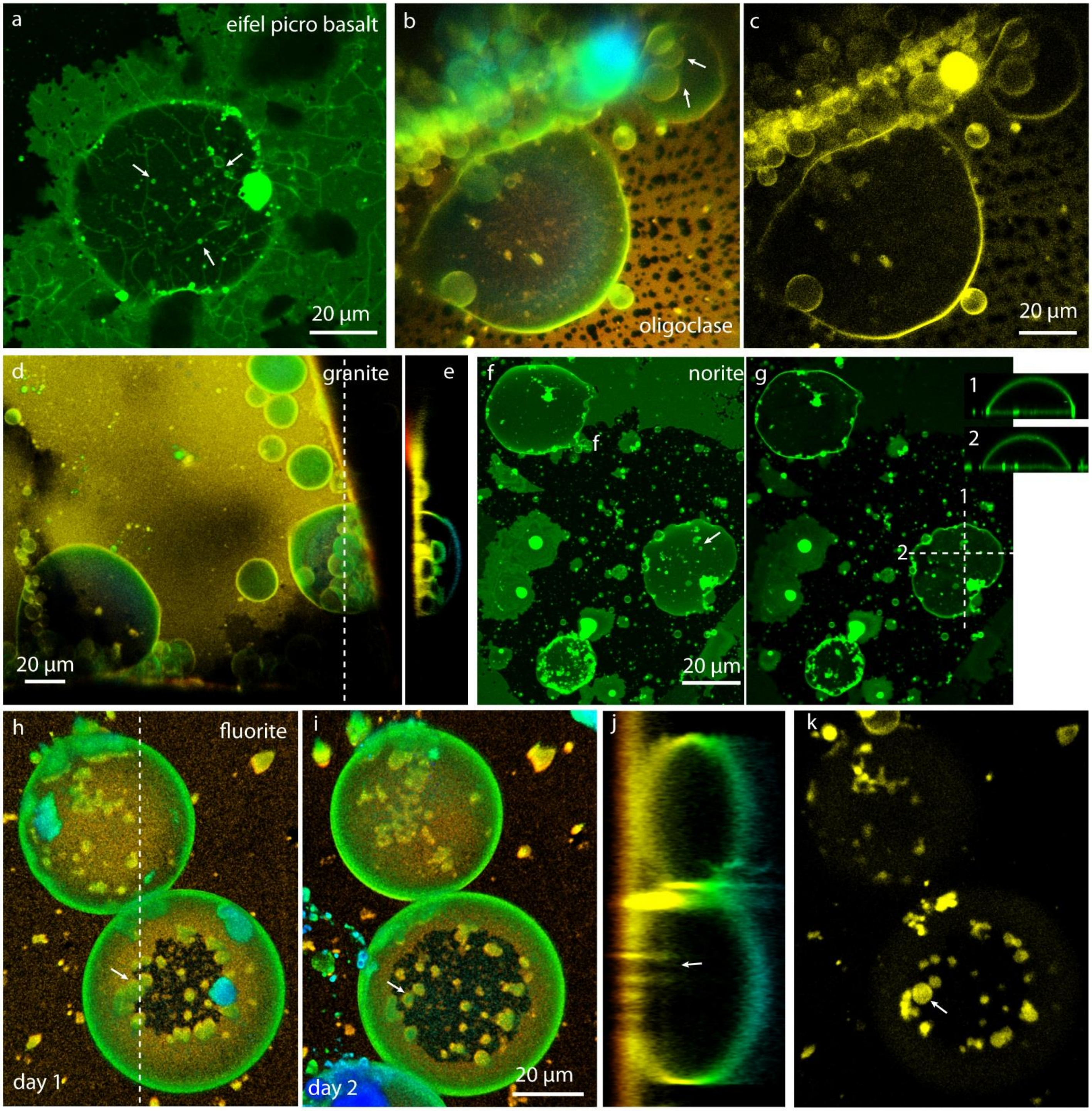


**Figure S5. Subcompartmentalization of protocells.** Confocal micrographs showing dome-shaped subcompartmentalized protocells on rock and mineral surfaces: **(a)** eifel picro-basalt, **(b-c)** oligoclase, **(d-e)** granite, **(f-g)** norite, **(h-j)** fluorite. (b), (d), (f) and (h-i) are epi-fluorescence projections of protocells. Panels (c), (g) and (k) show cross sections in (x-y plane) of (b), (f) and (h), respectively. (e), numbered insets in (g) and (j) show cross sections along the dashed lines in (d), (g) and (h), respectively (x-z plane). White arrows in (a), (b), (f) and (h-k) point to subcompartments.

### S7. SEM-EDX analysis

#### S7.1. Olivine

SEM-EDX analysis of olivine which enhances protocell formation on several surface regions, have been performed, and shown in **Fig. S6**. Results revealed olivine with composition of $Mg_2SiO_4$ and $Fe_2SiO_4$.

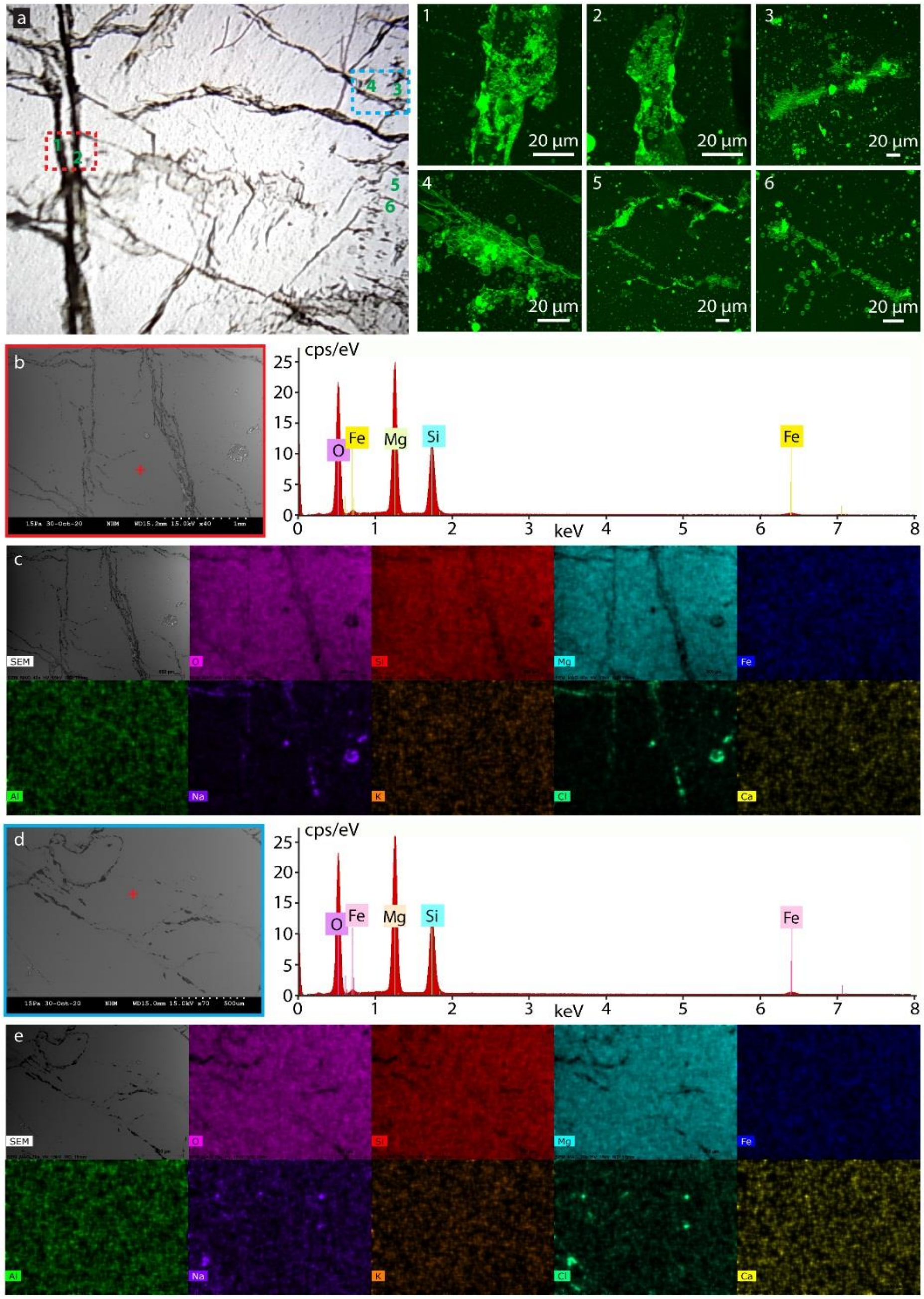


**Figure S6. SEM-EDX analysis of surface of olivine. (a)** Photograph of olivine surface. Regions that lipid nanotube-vesicle networks and protocell colonies on adhered bilayer are numbered 1-6 and confocal micrographs of corresponding regions are shown to the left. **(b)** SEM image of the region marked with red dashed frame in (a) and SEM-EDX point analysis plot. **(c)** SEM-EDX scans of the region in (b) showing different elements. **(d)** SEM image of the region marked with blue dashed frame in (a) and SEM-EDX point analysis plot. **(e)** SEM-EDX scans of the region in (d) showing different elements.

#### S7.2. Fluorite

SEM-EDX analysis of fluorite which enhances protocell formation on several surface regions, have been performed, and shown in **Fig. S8**. Results revealed pure fluorite with composition of $CaF_2$.

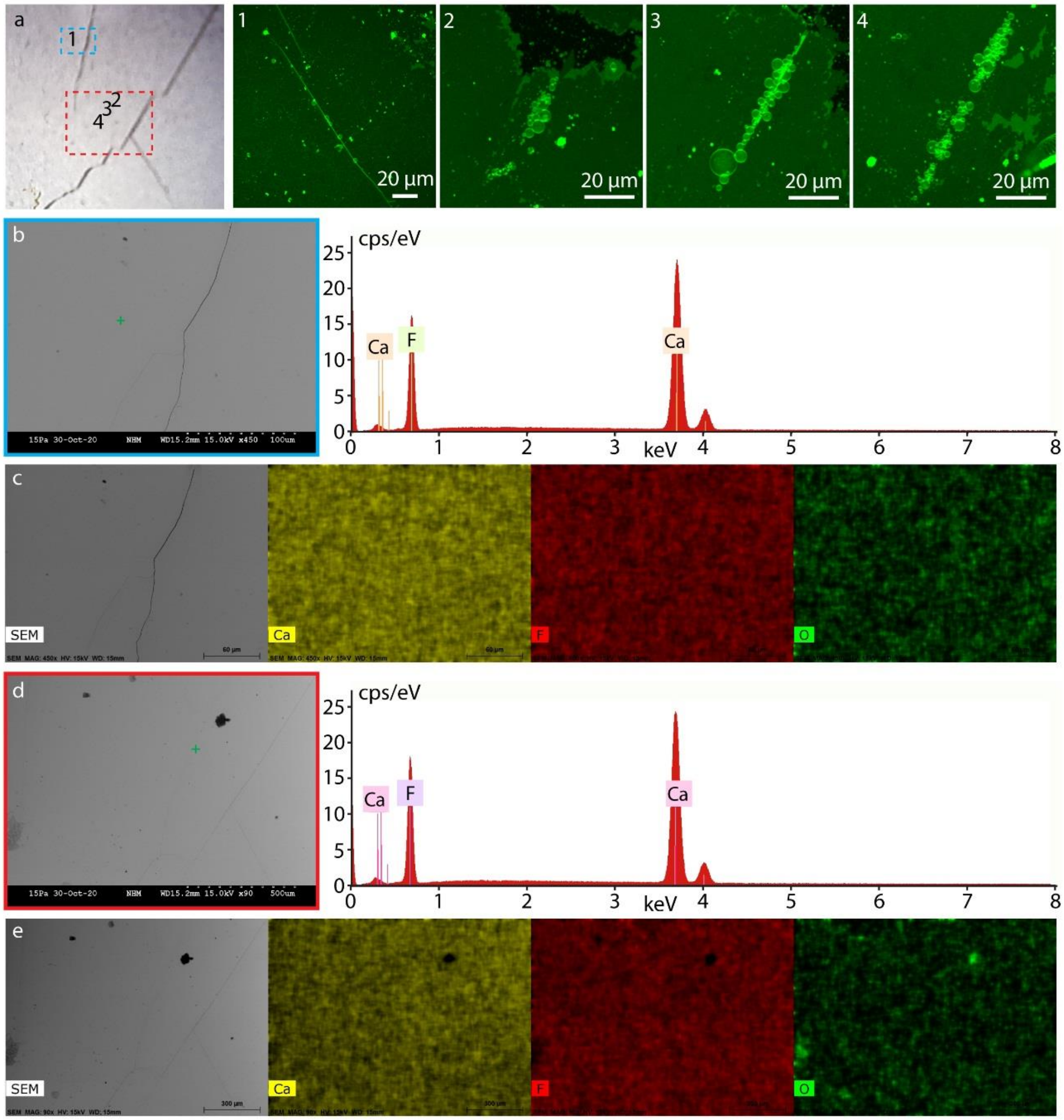


**Figure S7. SEM-EDX analysis of surface of fluorite. (a)** Photograph of fluorite surface. Regions that lipid nanotube-vesicle networks and protocell colonies on adhered bilayer are numbered 1-4 and confocal micrographs of corresponding regions are shown to the left. **(b)** SEM image of the region marked with blue dashed frame in (a) and SEM-EDX point analysis plot. **(c)** SEM-EDX scans of the region in (b) showing different elements. **(d)** SEM image of the region marked with red dashed frame in (a) and SEM-EDX point analysis plot. **(e)** SEM-EDX scans of the region in (d) showing different elements.

#### S7.3. Olivine-rich lava rock

SEM-EDX analysis of olivine-rich lava rock which enhances protocell formation on several surface regions, have been performed, and shown in **Fig. S8**. Results revealed olivine with composition of $Mg_2SiO_4$, $Fe_2SiO_4$ and $Ca_2SiO_4$ in the regions amplifying protocell formation.

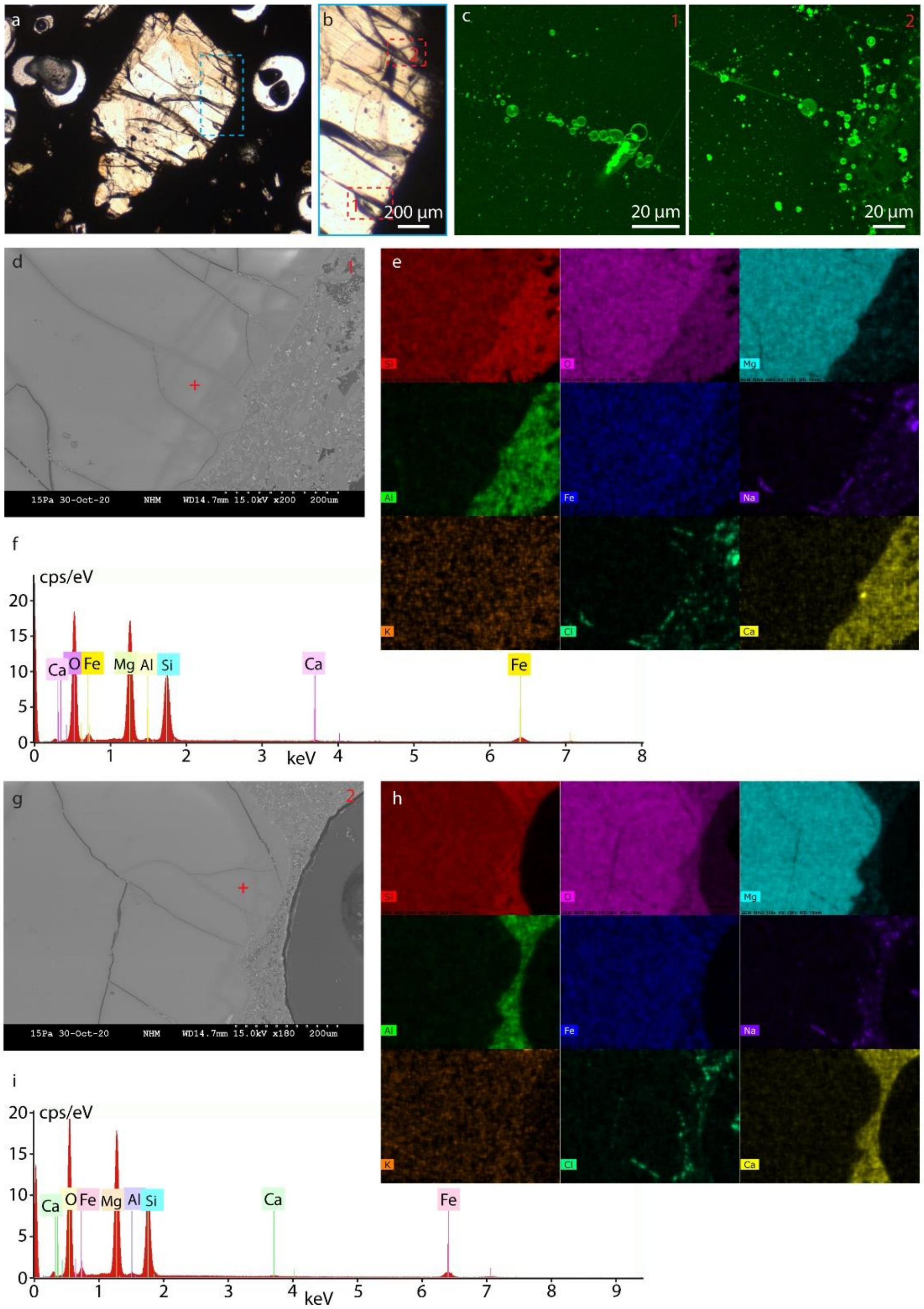


**Figure S8. SEM-EDX analysis of surface of olivine-rich lava rock. (a-b)** Transmission image of olivine-rich lava rock. (b) is the region with blue dashed frame in (a). **(c)** Confocal micrographs showing protocell formation corresponding to the numbered areas in (b). **(d)** and **(g)** are SEM images of the 1st and 2nd regions marked with red dashed frame in (b). **(e)** and **(h)** are SEM-EDX scans of the regions in (d) and (g), respectively showing different elements. **(f)** and **(i)** show SEM-EDX point analysis plots of the marked points in (d) and (g), respectively.

#### S7.4. Eclogite

SEM-EDX analysis of eclogite containing different grains/minerals is provided in **Fig. S9**. Results revealed that protocell amplification occurs at the interface of quartz-containing grains.

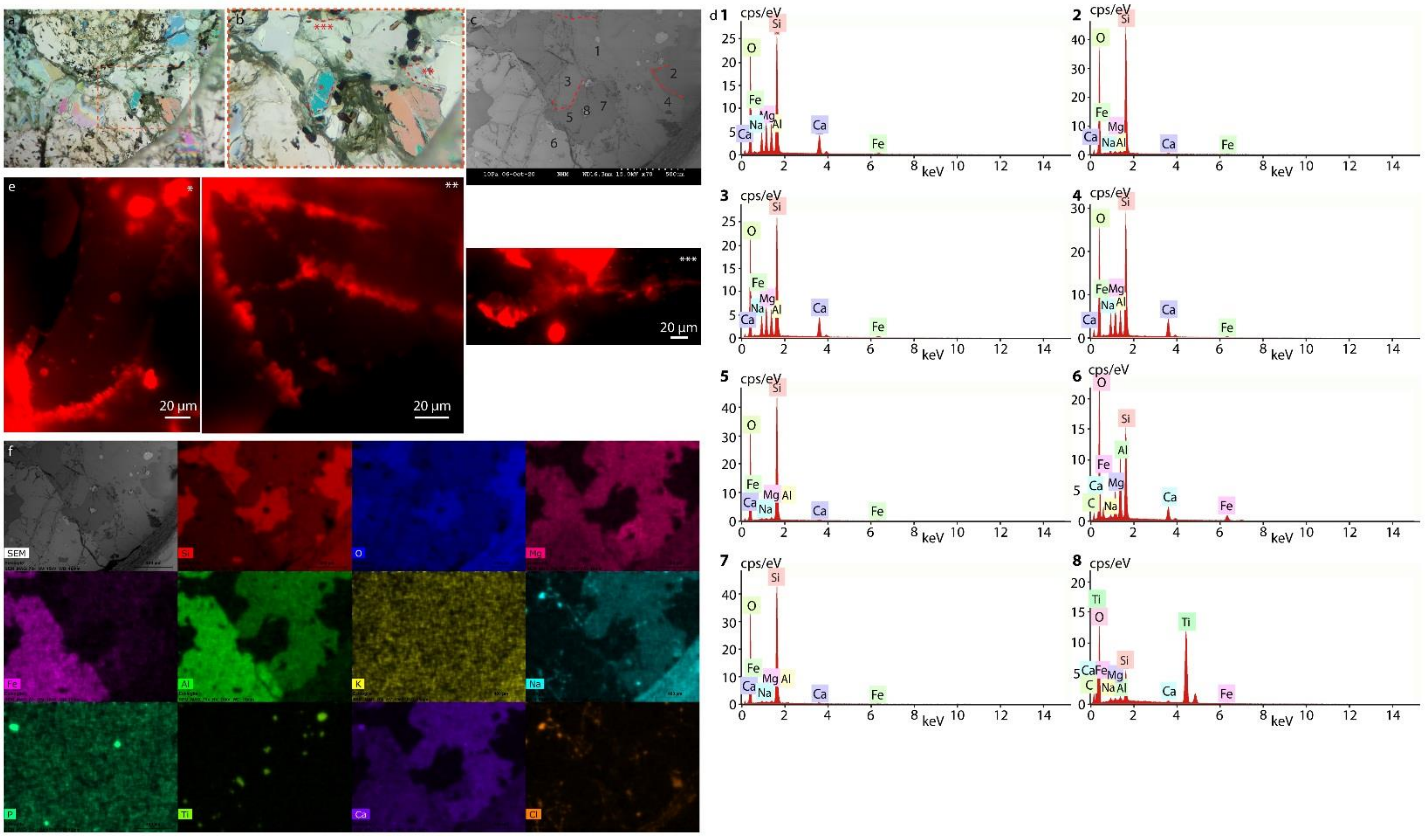


**Figure S9. SEM-EDX analysis of surface of eclogite rock. (a-b)** Polarized microscopy image of eclogite showing different domains. (b) is the region with red dashed frame in (a). **(c)** SEM image of the region in (b). **(d)** Plots of SEM-EDX point analysis on numbered points in (c). **(e)** Confocal micrographs showing protocell formation corresponding to the areas marked with different number of stars. Shape of the grain boundary where protocells are formed are also drawn with dashed lines in (b). **(f)** shows the SEM-EDX scans of the region in (b) and (c) showing different elements.

### S7.5. Granite

SEM-EDX analysis of granite rock containing different grains/minerals is provided in **Fig. S10**. Results revealed that protocell amplification is enhanced at the interface of quartz-containing grains.

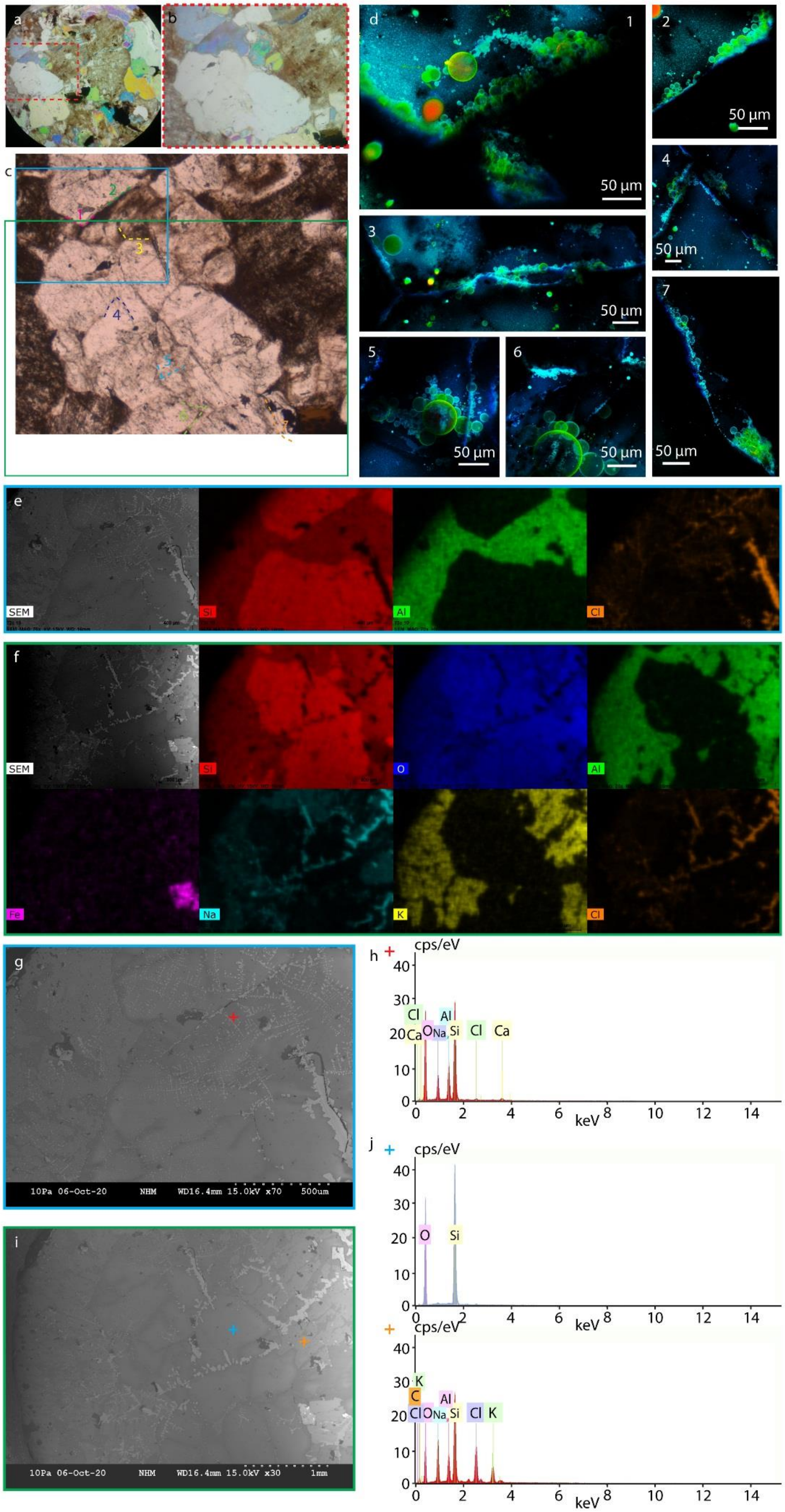
a
b
c
d
1
2
3
4
5
6
7
50 µm
e
SEM
Si
Al
Cl
f
SEM
Si
O
Al
Fe
Na
K
Cl
g
10Pa 06-Oct-20 NHM WD16.4mm 15.0kV x70 500um
h
cps/eV
keV
O Na Al Si Cl Ca
i
10Pa 06-Oct-20 NHM WD16.4mm 15.0kV x30 1mm
j
cps/eV
keV
O Si
C K Cl O Na Al Si Cl K

**Figure S10. SEM-EDX analysis of granite surface. (a-b)** Polarized microscopy image of granite showing different domains/minerals. (b) is the region with red dashed frame in (a). **(c)** Transmission image of the region shown in (b). **(d)** Confocal micrographs showing protocell formation corresponding to the areas numbered 1-7 in (c). Shape of the grain boundaries where protocells formed are also drawn with dashed lines in (c). **(e-f)** show the SEM-EDX scans and **(g)** and **(i)** show the SEM image of the regions defined with blue and green frames in (c), respectively. **(h)** and **(j)** are the plots of SEM-EDX point analysis of marked points in (g) and (i).

#### S7.6. Martian meteorite NWA7533

SEM-EDX analysis has been performed on several surface regions of Martian meteorite NWA 7533. While results, summarized in **Fig. S11-S14,** did not reveal any preference for specific element and amplification of protocells, results in **Fig. S15-S16** indicated consistent formation of protocells on plagioclase mineral. Additionally, analysis on ruptured areas with nanotubular structures shown in **Fig. S17-S18** did not point to a specific mineral.

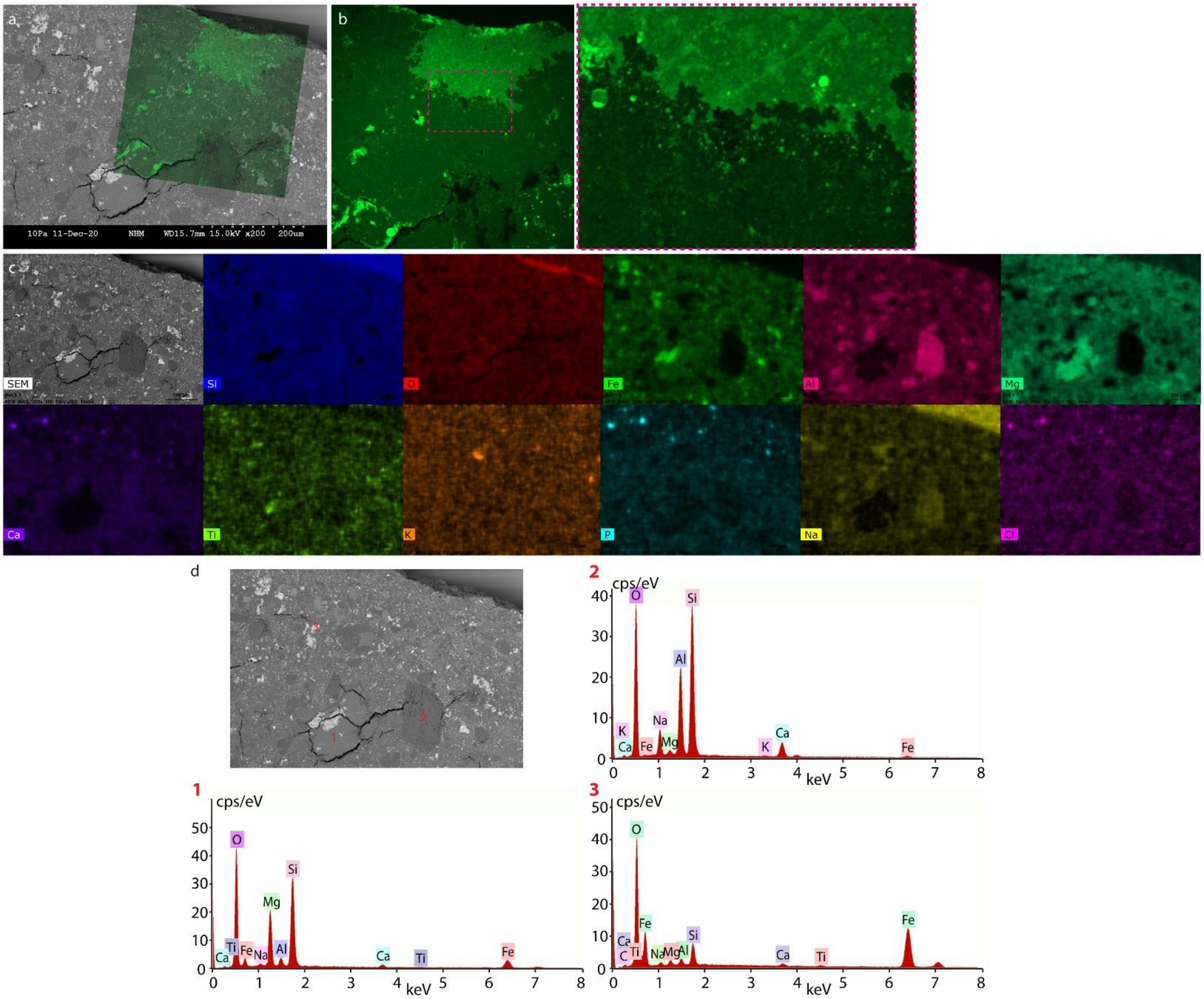


**Figure S11. SEM-EDX analysis of a region on the surface of NWA7533 meteorite amplifying protocell formation. (a)** Superimposed SEM image of meteorite and positioned fluorescence micrograph of the membrane. **(b)** Confocal micrograph showing a double bilayer dewetting a single bilayer, formed nanotubular network and small, nanotube-attached vesicles adhered on the bilayer shown in **Fig. 4e**. **(c)** SEM-EDX scans of the region in (a) showing different elements. **(d)** SEM image showing three regions that SEM-EDX point analysis were performed and corresponding analysis plots.

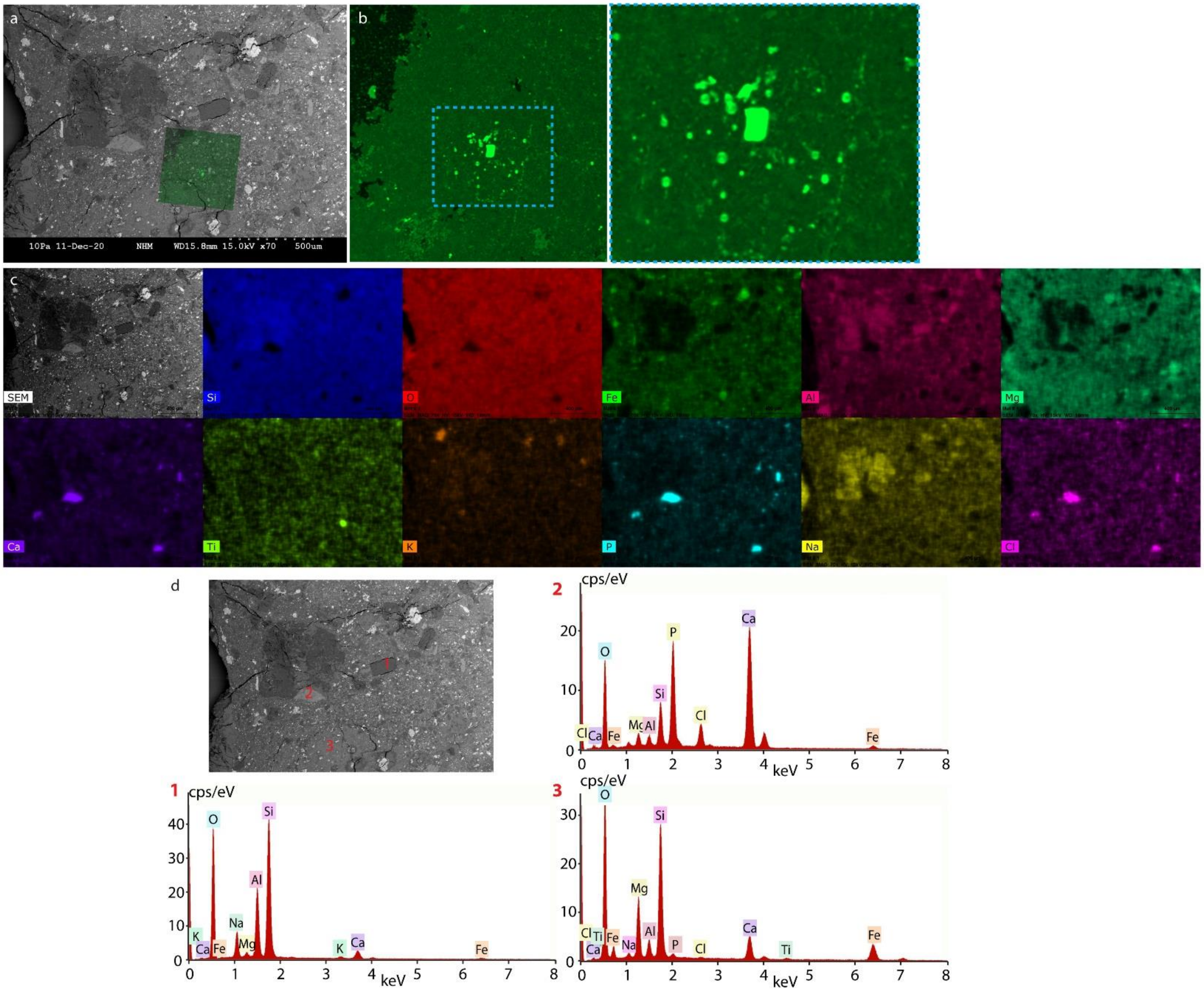


**Figure S12. SEM-EDX analysis of a region on the surface of NWA7533 meteorite amplifying protocell formation. (a)** Superimposed SEM image of meteorite and positioned fluorescence micrograph of the membrane. **(b)** Confocal micrograph showing formation of small protocells from short nanotubes on a bilayer. **(c)** SEM-EDX scans of the region in (a) showing different elements. **(d)** SEM image showing three regions that SEM-EDX point analysis were performed and corresponding analysis plots.

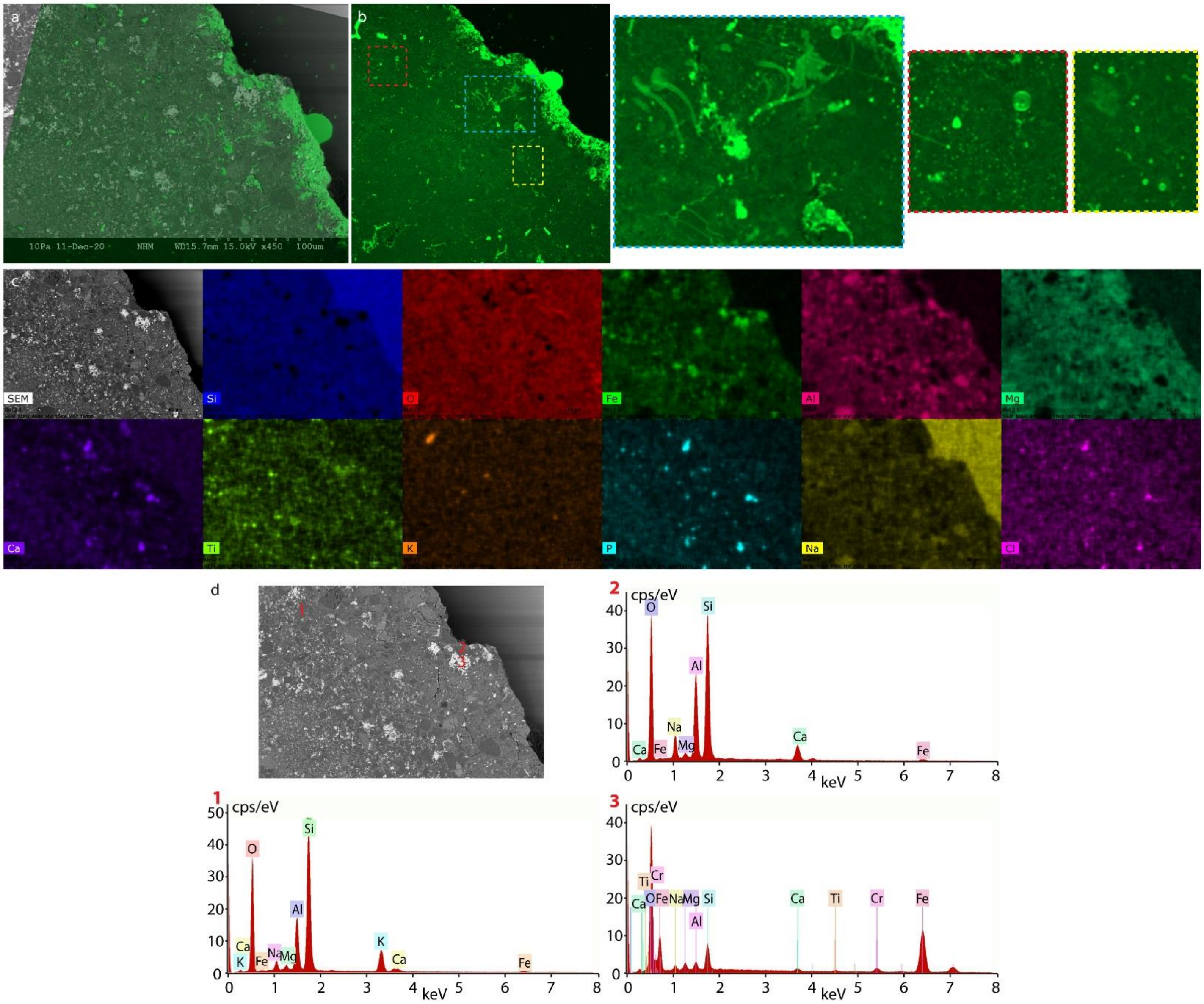


**Figure S13. SEM-EDX analysis of a region on the surface of NWA7533 meteorite amplifying protocell formation. (a)** Superimposed SEM image of meteorite and positioned fluorescence micrograph of the membrane. **(b)** Confocal micrograph showing short nanotubes and small protocells on a bilayer. **(c)** SEM-EDX scans of the region in (a) showing different elements. **(d)** SEM image showing three regions that SEM-EDX point analysis were performed and corresponding analysis plots.

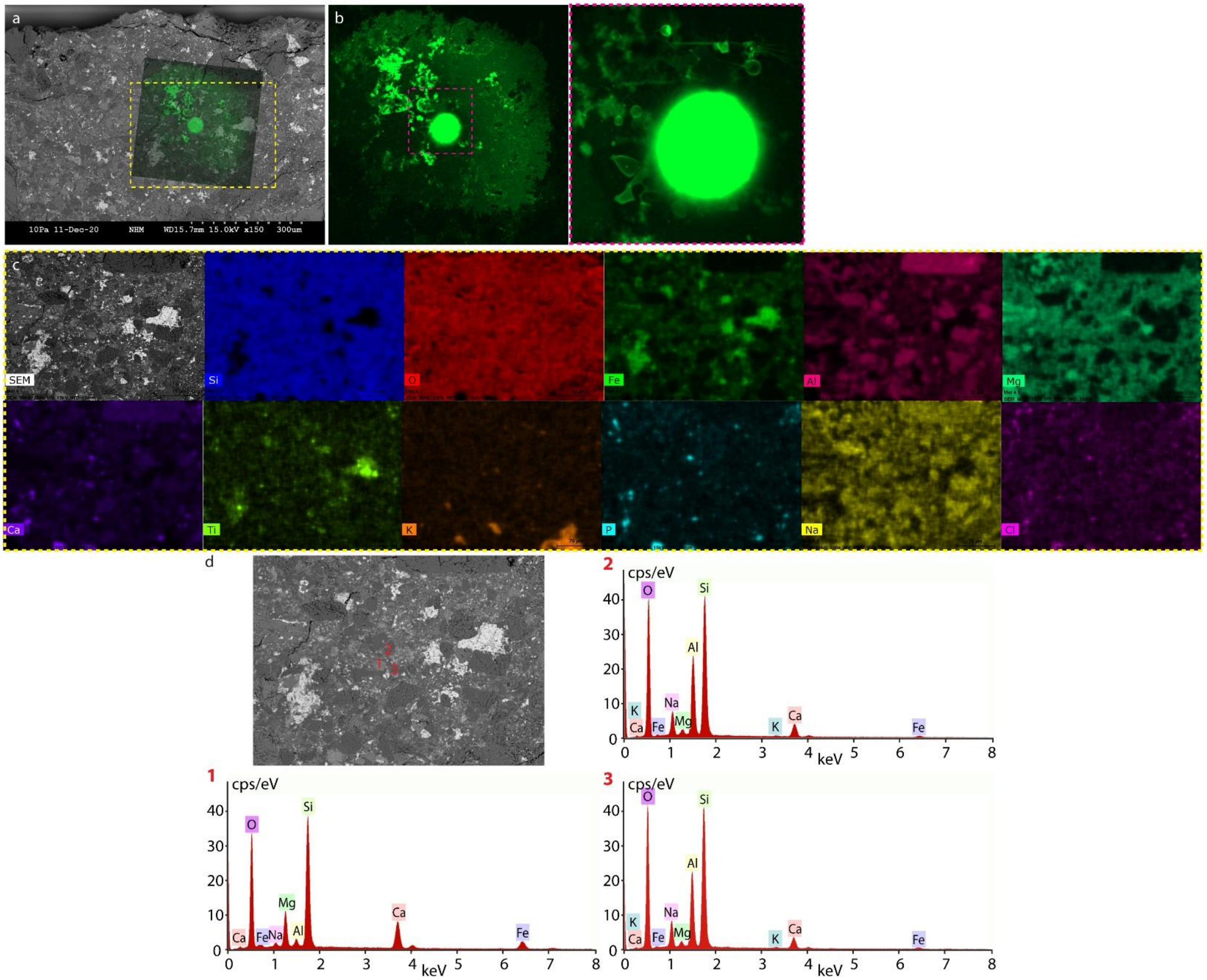


**Figure S14. SEM-EDX analysis of a region on the surface of NWA7533 meteorite amplifying protocell formation.** **(a)** Superimposed SEM image of meteorite and positioned fluorescence micrograph of the membrane. **(b)** Confocal micrograph showing a multilamellar lipid reservoir which circularly spread to form membrane patch having lipid nanotubes and protocells. **(c)** SEM-EDX scans of the region shown with yellow dashed frame in (a) showing different elements. **(d)** SEM image showing three regions that SEM-EDX point analysis were performed and corresponding analysis plots.

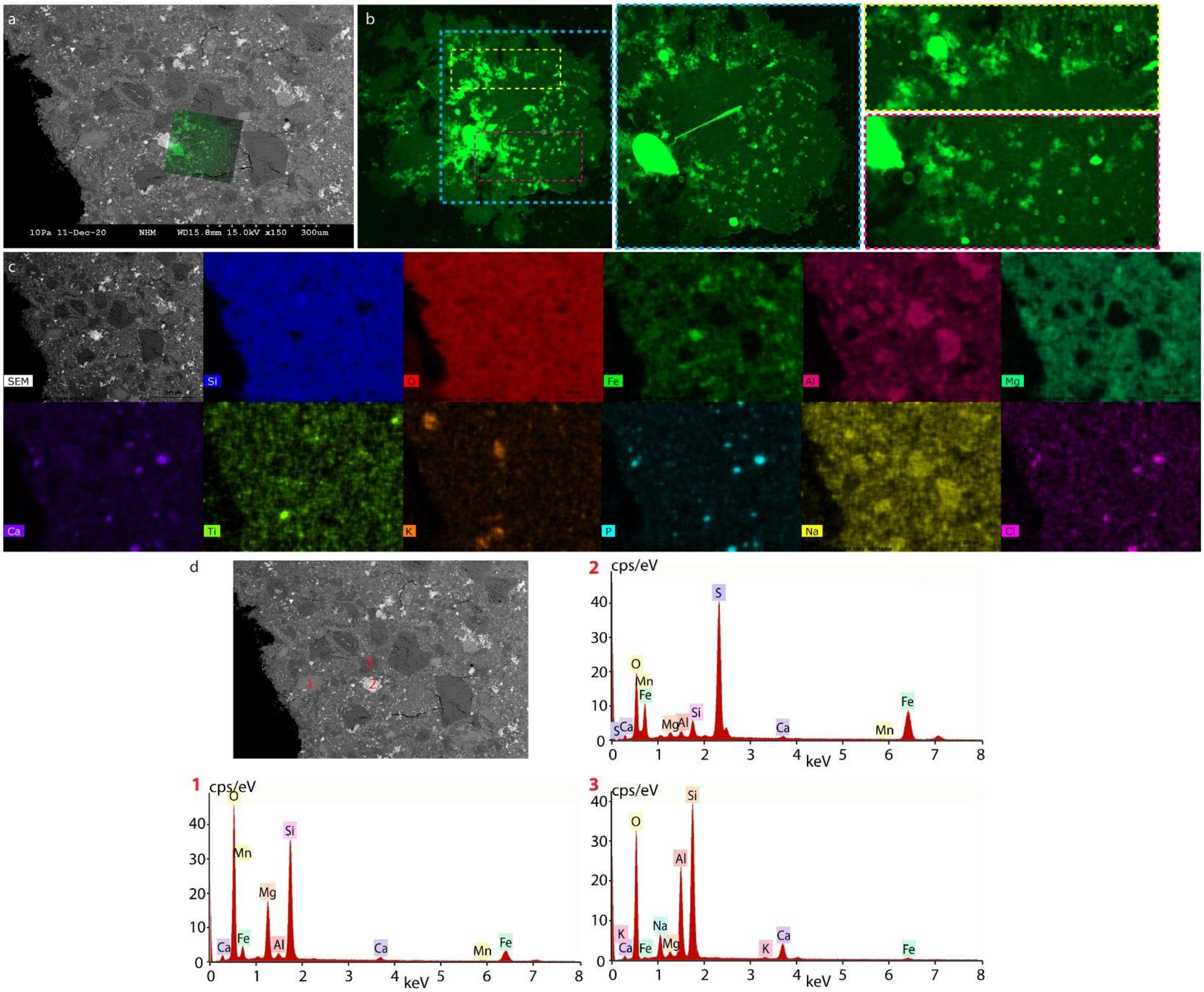


**Figure S15. SEM-EDX analysis of a region on the surface of NWA7533 meteorite amplifying protocell formation. (a)** Superimposed SEM image of meteorite and positioned fluorescence micrograph of the membrane. **(b)** Confocal micrograph showing a multilamellar lipid reservoir which circularly spread to form membrane patch and grown protocells around it. **(c)** SEM-EDX scans of the region in (a) showing different elements. **(d)** SEM image showing three regions that SEM-EDX point analysis were performed and corresponding analysis plots.

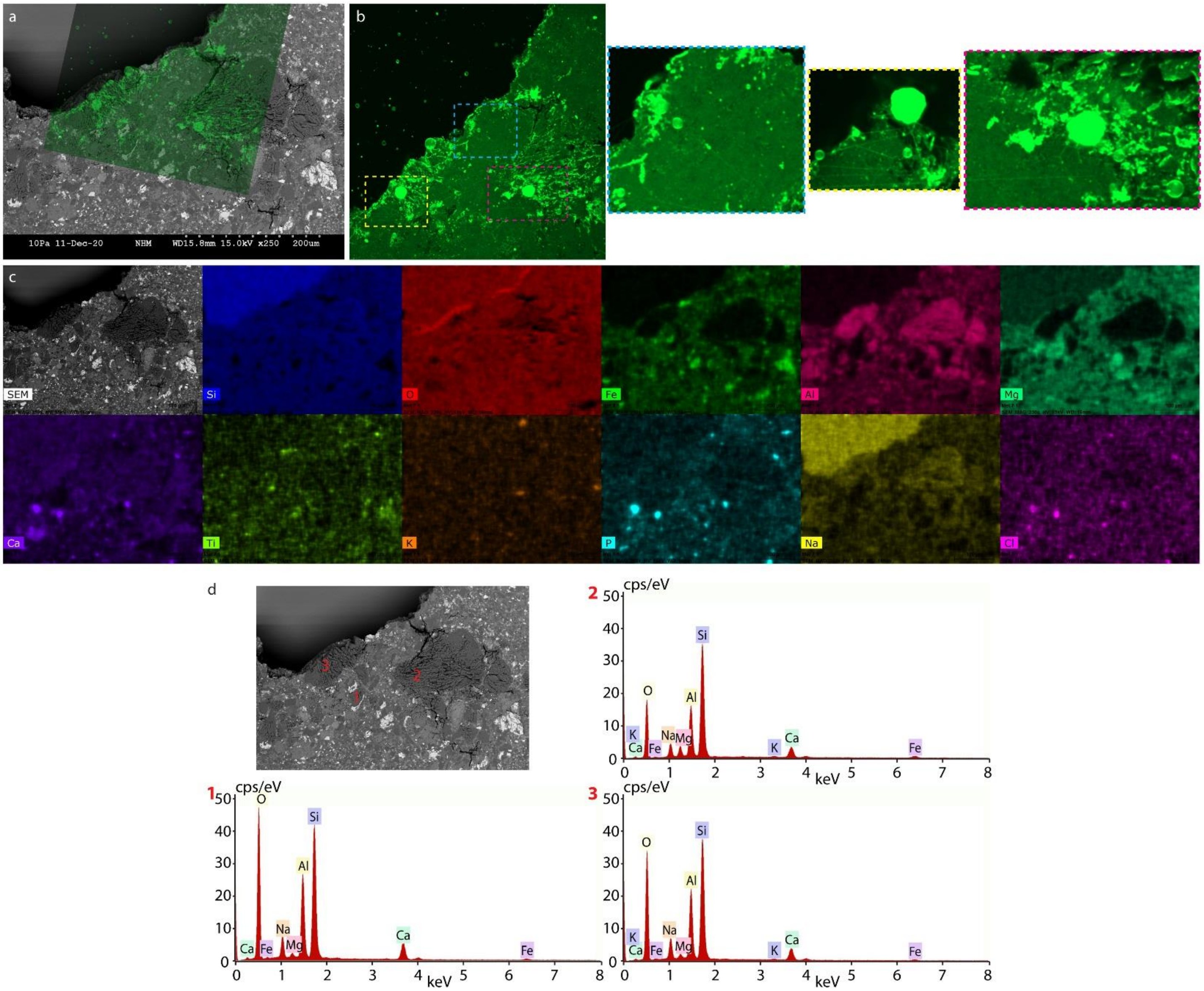


**Figure S16. SEM-EDX analysis of a region on the surface of NWA7533 meteorite amplifying protocell formation. (a)** Superimposed SEM image of meteorite and positioned fluorescence micrograph of the membrane. **(b)** Confocal micrograph showing nanotubes and grown protocells on them. **(c)** SEM-EDX scans of the region in (a) showing different elements. **(d)** SEM image showing three regions that SEM-EDX point analysis were performed and corresponding analysis plots.

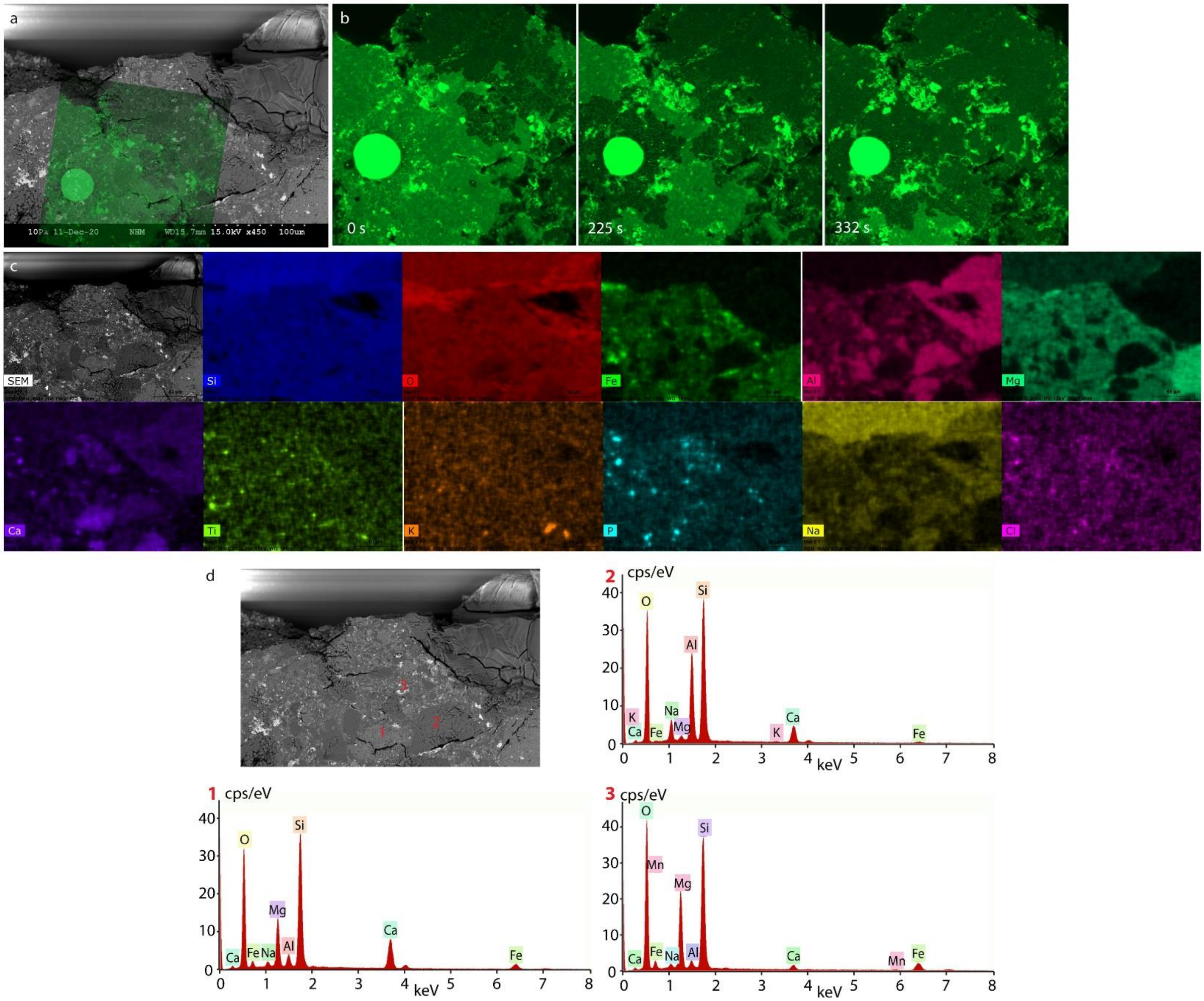


**Figure S17. SEM-EDX analysis of a region on the surface of NWA7533 meteorite amplifying protocell formation. (a)** Superimposed SEM image of meteorite and positioned fluorescence micrograph of the membrane. **(b)** Confocal micrograph time series showing double bilayer dewetting a single bilayer and formed nanotubular network shown in **SI Movie 2** ‘Rupture 1’. **(c)** SEM-EDX scans of the region in (a) showing different elements. **(d)** SEM image showing three regions that SEM-EDX point analysis were performed and corresponding analysis plots.

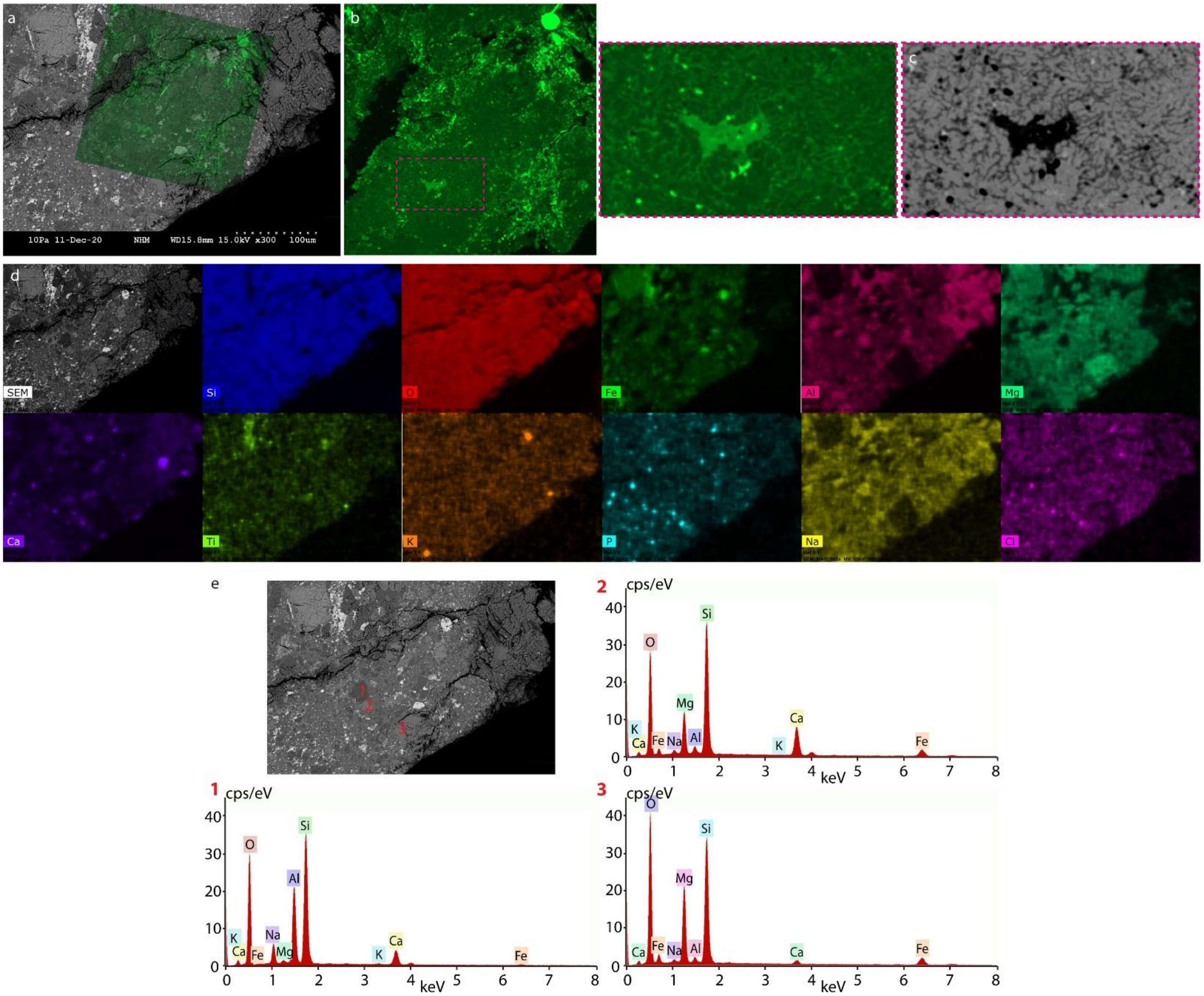


**Figure S18. SEM-EDX analysis of a region on the surface of NWA7533 meteorite amplifying protocell formation. (a)** Superimposed SEM image of meteorite and positioned fluorescence micrograph of the membrane. **(b)** Confocal micrograph showing a network of lipid nanotubes and small emerging protocells on them. **(c)** inverted confocal micrograph of a region shown in (b) with nanotubes. **(d)** SEM-EDX scans of the region in (a) showing different elements. **(e)** SEM image showing three regions that SEM-EDX point analysis were performed and corresponding analysis plots.

### S8. X-ray fluorescence (XRF) spectroscopy and inductively coupled plasma mass spectrometry (ICP-MS) analysis

Result of whole rock analysis with XRF spectroscopy on some of the surfaces is shown in **Fig. S22**. Our observations did not indicate any correlation between protocell formation and presence of composition of these surfaces.

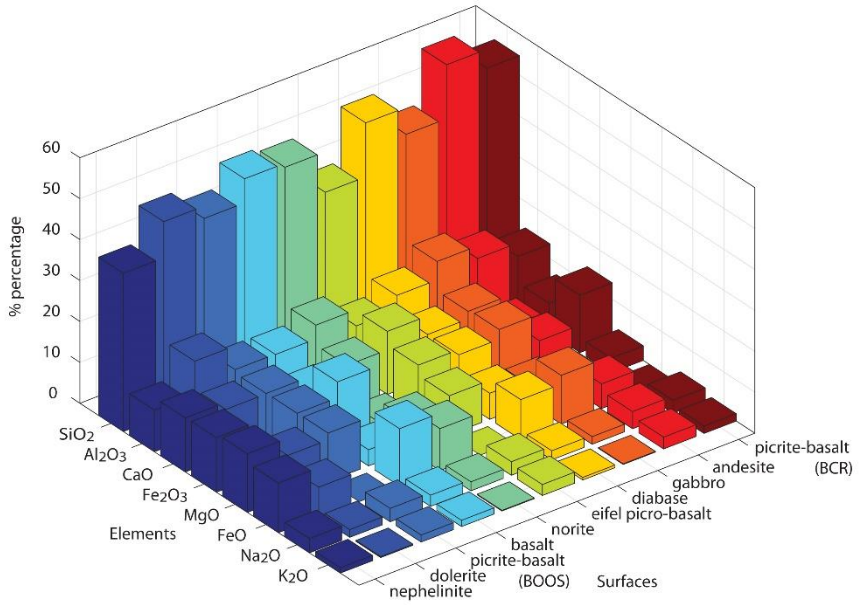


**Figure S19. 3D bar plot showing major elemental composition of glasses determined with XRF spectroscopy.**

### S9. Extended Figure 4: Protocell formation on the NWA 7533

**Fig. S20a-e** shows more examples of protocell formation and foam-like colony formation on Martian meteorite NWA 7533 (**Fig. 4a-e**). Additional examples to development of smaller colonies from foam-like structures shown in **Fig. 4b** are also provided in **Fig. S20d-e**. Both floral and fractal rupture morphologies (pink arrows in **Fig. S20g-i**) are observed on the meteorite. Fluorescence intensity profiles along the dashed lines in **Fig. S20f-i** show 2x the intensity in double bilayer regions (bright green) compared to the single bilayer regions (dark green). Whole process of membrane rupturing on the meteorite is shown in **Movie S2** in which the region shown in **Fig. S20i** and **Fig. S20f** corresponds to ‘Rupture 2 and 3’**,** respectively. Intermittent behavior of fractal rupturing can be observed in the ‘Rupture 2’ of **Movie S2**.

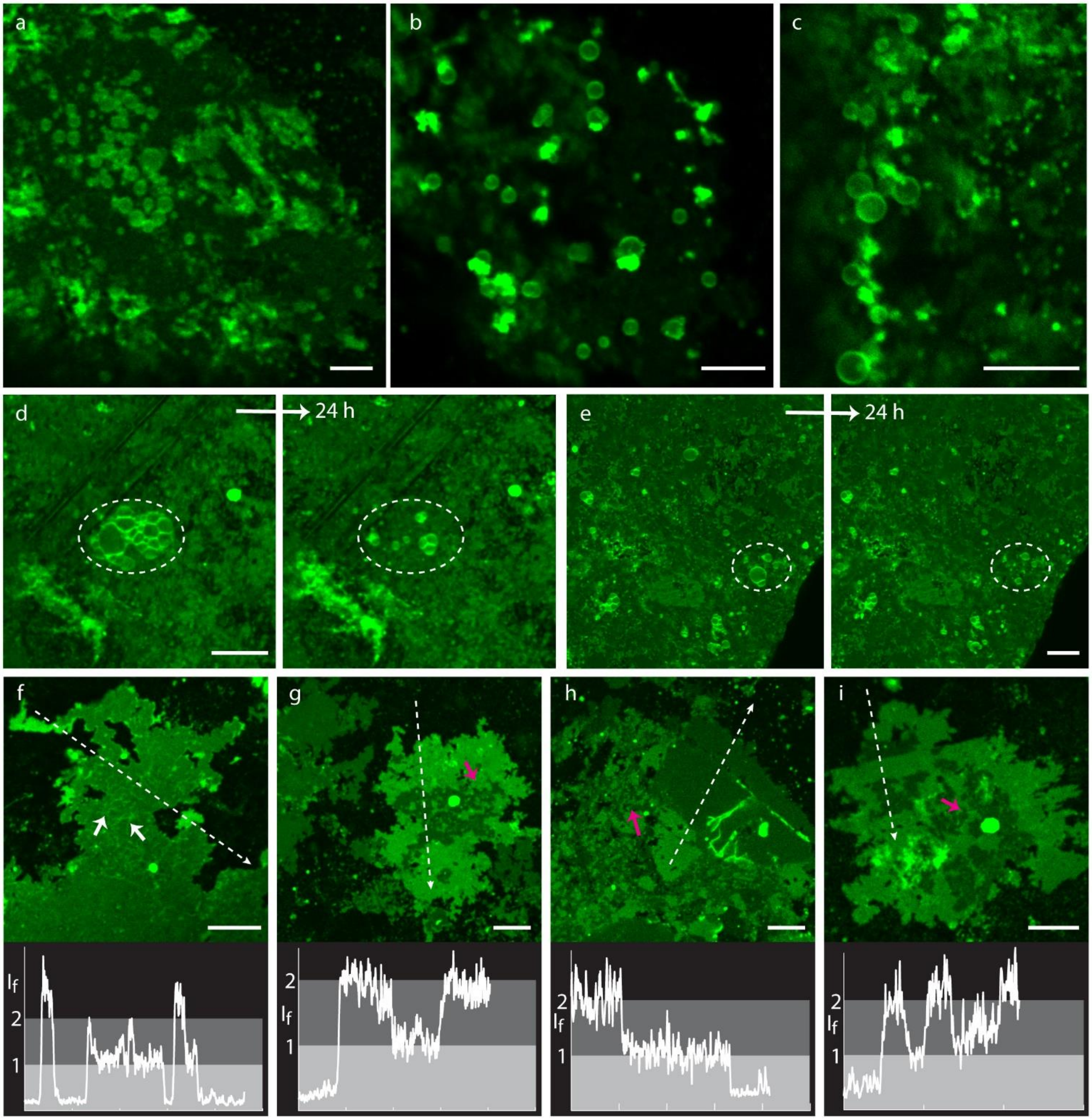


**Figure S20. Formation of protocells on the Martian meteorite (NWA 7533) specimen.** Confocal micrographs showing: protocell formation on lipid patches adhered onto the meteorite **(a-c)**, division of the colonies after a day to smaller units **(d-e)**. The position of the original structures and their corresponding daughter cell structures have been encircled with white dashed lines. **(f-i)** Confocal micrographs showing double bilayers (bright green) and single bilayers (dark green). The plots under every micrograph shows the fluorescence intensity profile over the dashed lines in the micrograph above. Pink arrows in **(g-i)** show fractal type of ruptures on membranes. White arrows in **(f)** show the nanotubes residing on a bilayer. Scale bars: 10 µm.

### S10. Extended Figure 5: RNA and DNA encapsulation

Another example of RNA encapsulation (complementary to **Fig. 5a-d**) is provided in **Fig. S21**. Three additional examples to DNA encapsulation and DNA reactions (complementary to **Fig. 5f-j**) are provided in **Fig. S22**.

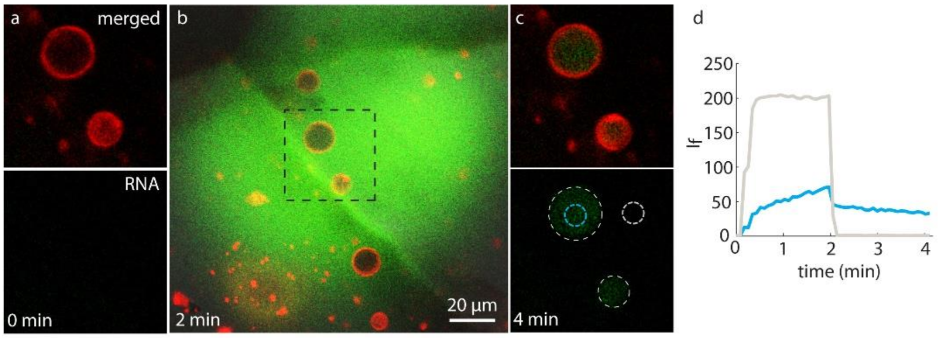


**Figure S21. RNA encapsulation. (a-c)** confocal micrographs showing protocells on eclogite before **(a)**, during **(b)**, and after **(c)** RNA (green fluorescence) superfusion. **(a)** and **(c)** represent the area framed in black dashed lines in **(b)**. **(d)** fluorescence intensity of the regions of interest in **(c)** (blue and gray dashed circles), over time. The vesicle on top takes up the RNA and maintains it over 4 minutes. The sharp drop in ambient RNA intensity (grey line) corresponds to the termination of controlled superfusion.

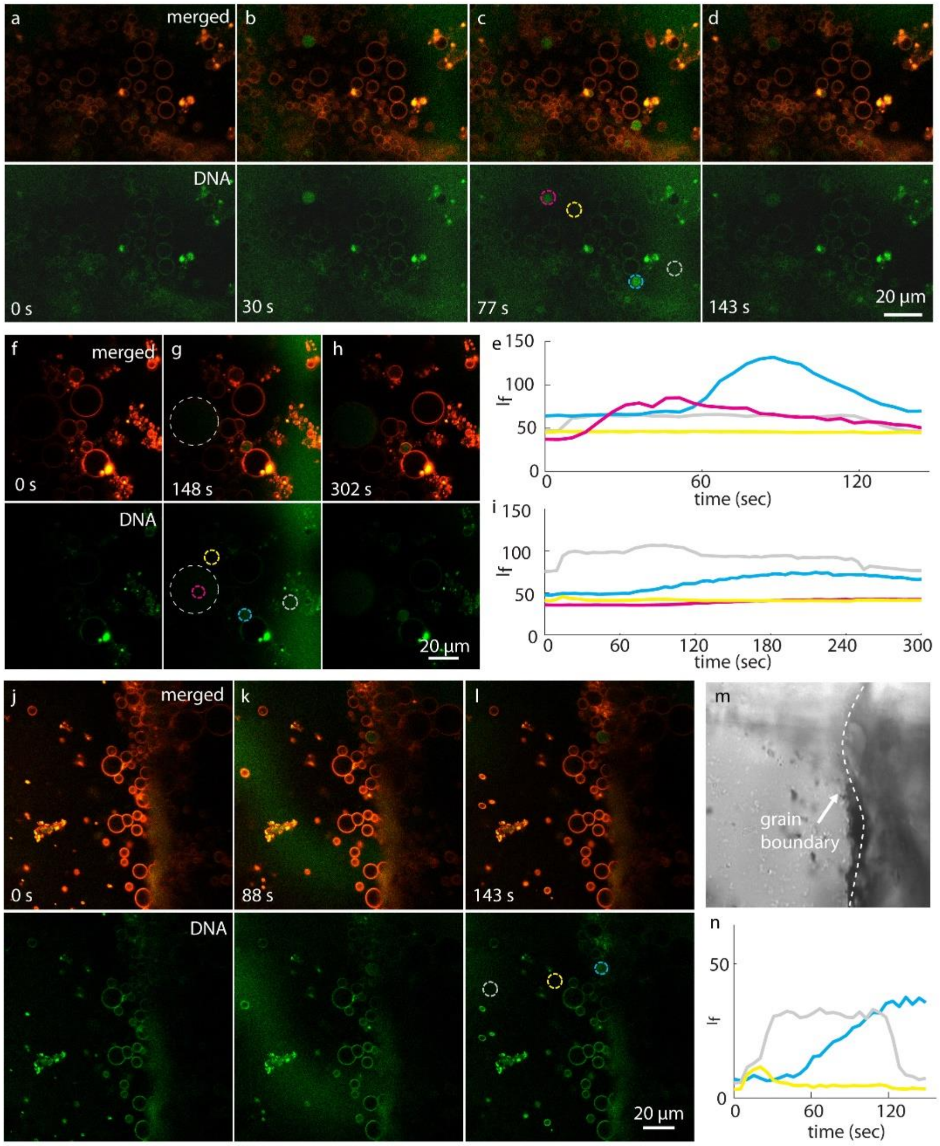


**Figure S22. DNA encapsulation and entropy driven DNA strand displacement reaction inside the protocells. (a-d)**, **(f-h)**, **(j-l)** confocal micrographs showing the encapsulation of ssDNA resulting in DNA strand replacement inside the protocells adhered on granite. The entry of the ssDNA leads to melting of annealed dsDNA previously encapsulated inside the protocells (red fluorescence) and hybridization of one of the strands with the newly entering ssDNA (strand displacement). The released ssDNA becomes no longer quenched and starts to fluoresce (green fluorescence). **(e)**, **(i)** and **(n)** fluorescence intensity inside the ROIs circled in (c), (g), (l), respectively versus time. **(m)** transmission image showing the grain boundary of the region shown in **(j-l)**.

### S11. Heat induced protocell formation

We previously reported heat-induced protocell formation from lipid-nanotube network on nano-engineered silica substrates[4] and reproduced these experiments for this study (**Fig. S23a-c**). Similar rapid protocell formation is observed on natural quartz (**Fig. S23d-g**) using IR-B laser radiation through a flat optical fiber tip as described before. A 3.4 W λ=1470 nm semiconductor diode laser (Seminex Inc., USA) in combination with a 50 µm core diameter, 0.22 NA multimode optical fiber (Ocean Optics Inc., USA) were used. The fiber was positioned using a 3-axis water hydraulic micromanipulator (Narishige, Japan) and the tip was located at 50 µm from the surface, resulting in a volume of approximately 1 nL being efficiently heated. The laser current was adjusted for growth of protocells in temperatures reaching up to ~70 °C as determined directly by an in-situ micro-thermocouple[4].
Additionally, here we report formation of dome-shaped protocells either with the help of local temperature increase (**Fig. S23f-g**) or spontaneously at room temperature on surfaces like eclogite and calcite (**Fig. S23h-n**).

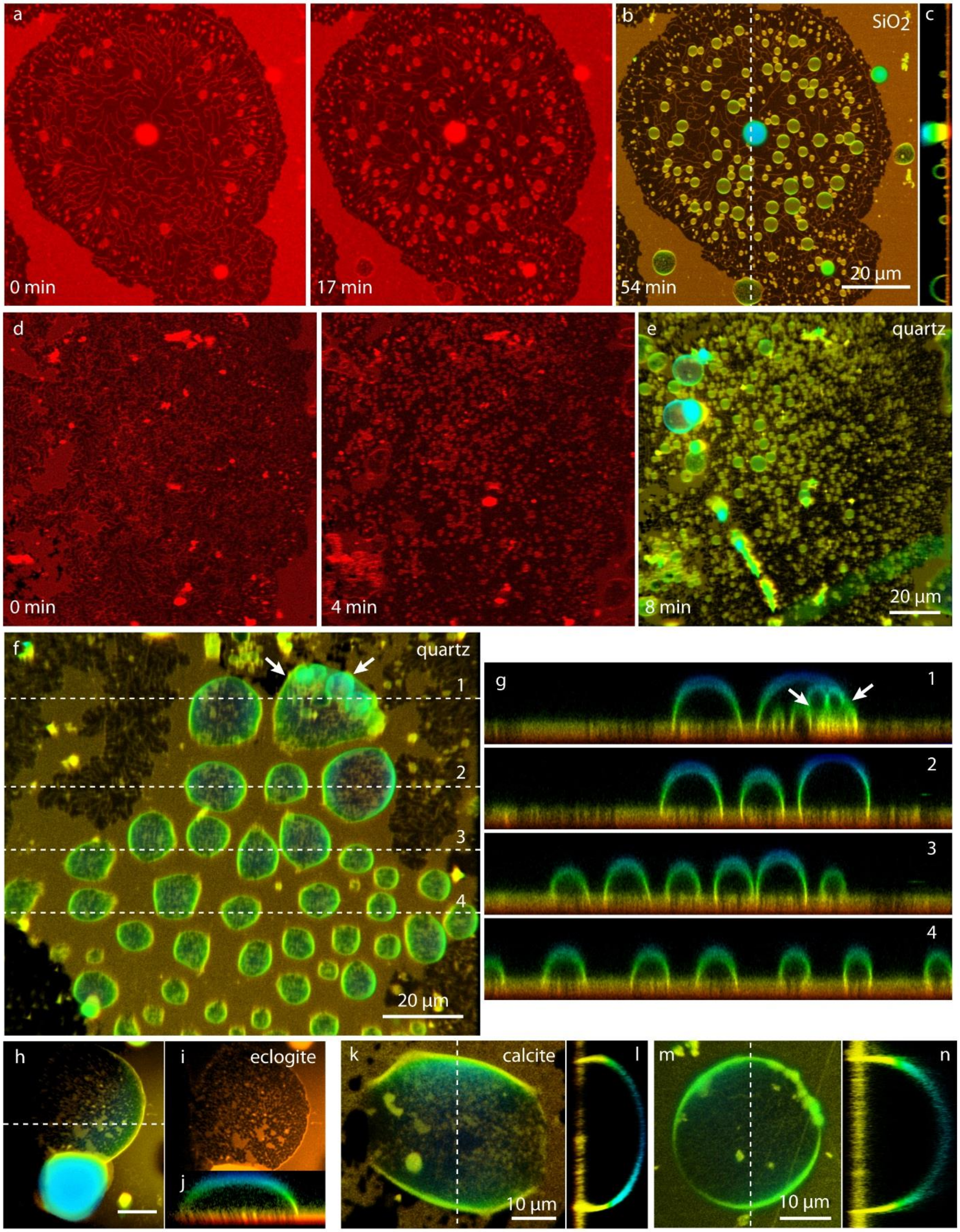


**Figure S23. Heat induced protocell formation.** Laser scanning confocal microscopy time series of heat-induced protocell formation from surface-adhered lipid nanotubes and formation of dome-shaped protocells from double bilayer on man-made $SiO_2$ **(a-c)** and quartz **(d-e)**. Panel (b) and (e) are epi-fluorescence projections of the rapidly grown protocells. (c) shows cross section along the dashed lines in (b) (x-z plane). **(f-g)** show another area with dome-shaped protocells and subcompartments formed from double bilayer on quartz surface. (f) is epifluorescence projection of protocells and (g) shows the cross sections along the numbered dashed lines in (f) (x-z plane). Arrows in (f-g) point to subcompartments. **(h-n)** show dome-shaped protocell formation from double bilayer on surfaces; (h-j) eclogite, (k-n) calcite at room temperature. (h), (k) and (m) are epifluorescence projections and (j), (l) and (n) are the cross sections along the dashed lines in these micrographs, respectively (x-z plane). (i) shows the cross section of (h) in x-y plane.

### S12. Supplementary movies

**Movie S1. RNA and DNA encapsulation. Movie S1** shows two examples of RNA encapsulation and five examples of DNA encapsulation and entropy driven DNA strand replacement reaction inside protocells. Movie is accelerated 15x. The region shown in 'RNA encapsulation 1-2' corresponds to **Fig. 5b-d** and **Fig. S21a-c**, respectively. The region shown in 'DNA reaction 1-5' corresponds to **Fig. 5h-j**, **Fig. 5g**, **Fig. S22a-d**, **Fig. 8j-l** and **Fig. S22f-h**, respectively.

**Movie S2. Membrane ruptures on the NWA7533. Movie S2** shows four examples of membrane ruptures on the Martian meteorite NWA7533. Movie is accelerated 15x. The region shown in 'Rupture 1-3' corresponds to **Fig. S17b**, **Fig. S20i** and **Fig. S20f**, respectively.
