## Supplementary material for "Spontaneous formation of prebiotic compartment colonies on Hadean Earth and pre-Noachian Mars": Materials and methods

***Preparation of thin rock, glass, mineral and meteorite sections.*** Petrographic thin sections of fluorite, eclogite, granite, oligoclase, quartz, olivine, calcite, olivine-rich lava rock, muscovite and marble were prepared by Vancouver Petrographics Ltd. (BC, Canada). Initially, thin sections were cut from a rock with a diamond saw. They were then mounted with an epoxy resin and polished on both sides optically flat to 170 µm thickness. The epoxy resin is acetone-soluble, so the thin sections can be recovered. For muscovite, a thin sheet was manually peeled from the surface gently using a scalpel.

The Northwest Africa 7533 meteorite was sectioned and a sample polished on both sides at the Facilities of the University of Copenhagen in a similar process to a thickness of 170 µm.

Thin sectioning (170 µm) of the dolerite, nephelinite, Eifel picro basalt, norite, diabase, picrite basalt, andesite, gabbro and basalt were performed at the Thin Section Lab at the Colorado School of Mines (CO, USA). The samples were cut into two pieces and polished afterwards. Complementary surfaces were polished on one side and embedded in an epoxy resin. One identical set of these surfaces was used to perform whole rock analyses using X-ray fluorescence spectroscopy (XRF). The analyses were performed and published by the US Geological Survey. The other set of samples was used for protocell formation experiments. All of the tested surfaces were rinsed with isopropanol, followed by rinsing with water and blow-drying with nitrogen. In case of re-use they were treated with oxygen plasma for 5 min before each experiment to eliminate any organic residues from the previous work.

**Synthesis of glasses**: The glasses were synthesized at the United States Geological Survey in Denver, Colorado USA as part of the Geological Reference Materials program[1]. Briefly, sufficient amount of base materials was pulverized and ground into a fine powder to prepare a reference material. Finely ground powders were mixed in homogenously. The mixture was melted in a high temperature furnace and subsequently flash-cooled to prevent crystallization. Formed glass-like material had known chemical composition and homogeneity at the 10-20 µm scale. The solid glass material was then cut into smaller pieces and sectioned as described above.

***Preparation of observation chamber.*** Thin mineral sections embedded in epoxy resin were submerged in acetone for 1-2 days until resin was completely dissolved. Sufficiently transparent minerals were gently glued onto a glass slide with a cm-sized circular perforation allowing contact with the objective lens of an inverted confocal microscope (Fig. 1). The hole in the glass slide was formed using a grinding pen with a sand paper circle nail drill (Cocraft, Clas Ohlson, Norway). Polydimethylsiloxane (PDMS) frame with dimensions 1.5 x 1.5 x 0.5 cm was adhered on top of the microscope slide to prepare an open top observation chamber[2]. Opaque minerals including the meteorite, were either gently glued on a 1 mm thick glass microscope slide or directly used as embedded in epoxy resin. PDMS frame was adhered on to the glass support they are embedded, to enable open top observation for the dip-in objectives of the upright confocal laser scanning microscope.

***Preparation of lipid reservoirs.*** A stock suspension of multilamellar lipid reservoirs containing 50% soybean polar lipid extract, 49% *E. coli* polar lipid extract  and 1% Rhodamine-PE or Carboxyfluorescein-PE (Avanti Polar Lipids, USA) were prepared by the dehydration/rehydration (gentle hydration) method as described before[2,3]. Briefly, lipids and lipid-conjugated fluorophores in designated ratios were mixed in chloroform reaching to a total concentration of 10 mg/ml. 300 µl of this solution was placed in a 10 ml round bottom flask and the chloroform was removed in a rotary evaporator at reduced pressure (20 kPa) over a period of 6 hours. The dry lipid film at the walls of the flask was rehydrated with 3 ml of PBS buffer containing 5 mM Trizma Base, 30 mM $K_3PO_4$, 30 mM $KH_2PO_4$, 3 mM $MgSO_4*7H_2O$ and 0.5 mM $Na_2EDTA$ and 30 µl glycerol. The pH was adjusted to 7.4 with $H_3PO_4$. The rehydrated lipid cake was kept at +4 °C overnight. In the final step, lipid cake was sonicated for 10 seconds at room temperature to induce the formation of giant vesicles of varying, mainly multiple lamellarity. Final stock suspension was aliquoted and kept at -20 °C till use.

Multilamellar lipid reservoirs were prepared from the stock lipid suspension kept at -20 °C. Briefly, 4 µl of stock lipid suspension was placed on a cover slip and dehydrated in an evacuated desiccator for 20 min. The dry lipid film was rehydrated with ~1 ml of HEPES buffer containing 10 mM HEPES and 100 mM NaCl (pH= 7.8, adjusted with NaOH) for 10 min to allow formation of MLVs. Using an automatic pipette 100-200 µl of MLV suspension was then transferred onto the prepared observation chamber with different mineral substrates containing ~1 ml of HEPES buffer with 10 mM HEPES, 100 mM NaCl and 4 mM $CaCl_2$ (pH= 7.8, adjusted with NaOH).

***Encapsulation of RNA with microfluidic pipette.*** An open-volume microfluidic pipette (Biopen, Fluicell AB, Sweden), positioned using a 3-axis water hydraulic micromanipulator (Narishige, Japan), was used to expose the surface-adhered matured protocells to 25 µM FAM-conjugated 10 base long poly-adenine RNA oligonucleotides (Dharmacon, USA) as described elsewhere[4].

***Encapsulation of DNA and strand displacement reaction.*** Three DNA oligonucleotides, purified by the manufacturer with HPLC (Integrated DNA technologies, USA), were used for the DNA reactions previously reported by Winfree et al.[5] The reactant sequences are presented below:

- reporter-ROX:/56-ROXN/CT TTC CTA CAC CTA CG
- reporter-RQ:TGG AGA CGT AGG TGT AGG AAA G/3IAbRQSp/
- output:CTT TCC TAC ACC TAC GTC TCC AAC TAA CTT ACG G

Thermal cycle was run using PTC-100 Programmable Thermal Controller (MJ Research, Inc., USA) to anneal 20 µM reporter-ROX and 20 µM reporter-RQ in TE buffer containing 10 mM Tris-HCl, 1mM EDTA and 12.5 mM $MgCl_2$ (pH= 8.0, adjusted with NaOH and HCl). The sample was incubated at 95 °C for 5 min followed by the cooling of the sample to 20 °C over 90 min. Subsequently, the sample was cooled down to +4 °C for storage until the encapsulation experiment. To encapsulate DNA dimers (reporter-ROX-reporter-RQ) inside the vesicles, 10 µM of DNA dimers in HEPES buffer containing 10 mM HEPES, 100 mM NaCl, 4 mM $CaCl_2$ was added to the ambient buffer of vesicles growing on the minerals. The excess dimers in the chamber were removed by gently exchanging the buffer with the dimer-free HEPES buffer using an automatic pipette in order to leave the dimers only inside the compartments. 10 µM output strand was superimposed locally with microfluidic pipette as described for RNA encapsulation and DNA reaction was observed thanks to generation of fluorescence signal from freed reporter-ROX strand.

***Microscopy imaging.*** An inverted confocal laser scanning microscopy system (Leica SP8, Germany), with an HCX PL APO CS 40x (NA 1.3) oil objective and 20x dry objective (NA X) was used for acquisition of the confocal images from transparent minerals. An upright confocal laser scanning microscopy system (BX81WI Olympus FluoView 1000, Japan), with 20x LumpPlanFl water immersion objective (NA 0.5) and 40x LumpPlanFl water immersion objective (NA 0.8) was used for opaque-samples. The utilized excitation/emission wavelengths for the imaging of the fluorophores, were as follows: $\lambda_{ex}$: 560 nm, $\lambda_{em}$: 583 nm for membrane fluorophore Rhodamine-PE, $\lambda_{ex}$: 488 nm, $\lambda_{em}$: 515 nm for Carboxyfluorescein-PE, $\lambda_{ex}$: 494 nm, $\lambda_{em}$: 525 nm for FAM, $\lambda_{ex}$: 588 nm, $\lambda_{em}$: 608 nm for ROX. Polarized microscope (Leica DMLP, Germany) equipped with Leica MC170HD camera and 2.5x objective were used for acquisition of polarization images of minerals.

***Surface microanalysis.*** Scanning electron microscopy (SEM) (S-3600N Hitachi, Japan) equipped with a 5-axis motorized stage with eucentric tilt (0 – 52°, depending on sample size) and 360° rotation was operated at variable pressure (VP) mode (10-200 Pa) without conductive coating of the mineral samples and meteorite for high resolution imaging. Semi- quantitative elemental analysis and hyperspectral mapping with <127 eV FWHM at MnKα energy resolution were done using EDX detector (Xflash® 5030 Bruker, USA) running on Quantax 400 (Esprit 1.9). XRF spectroscopy to determine major elemental composition were used on dolerite, nephelinite, Eifel picro basalt, picrite basalt, norite, diabase, picrite basalt, andesite, gabbro and basalt.

***Image processing/analysis.*** 3D fluorescence micrographs were reconstructed using the Leica Application Suite X Software (Leica Microsystems, Germany) and Imaris Viewer (Bitplane AG, Switzerland). Image enhancements to fluorescence micrographs were performed with the NIH Image-J Software and Adobe Photoshop CS4 (Adobe Systems, USA). The colors assigned to the labeled lipid membranes (red, green and depth coding) and to the labeled RNA and DNA oligonucleotides (green) are resulting from the false coloring of the gray scale images of the fluorescence signals, and are assigned arbitrarily. Schematic drawings and image overlays were created with Adobe Illustrator CS4 (Adobe Systems, USA). Fluorescence intensity profiles were drawn in Matlab R2018a. Fluorescence intensity measurements of different regions of interest on time series for RNA and DNA encapsulation experiments were done using the NIH Image-J Software and graphs were plotted in Matlab R2018a.

### Data availability

The data that support the findings of this study are available from the corresponding author upon reasonable request.

### Acknowledgements

We thank Lilian Garcia for providing the olivine-rich lava rock sample, Prof. Aldo Jesorka from Chalmers University of Technology, Sweden for his technical advice, Prof. Stephanie Warner and Dr. Agata Krzesinska from the University of Oslo for stimulating discussions. This work was made possible through financial support obtained from the Research Council of Norway (Forskningsrådet) Project Grant 274433, UiO: Life Sciences Convergence Environment, as well as the start up funding provided by the Centre for Molecular Medicine Norway (RCN 187615) & Faculty of Mathematics and Natural Sciences at the University of Oslo. I.P. greatly acknowledges the European Union's Horizon 2020 research and innovation programme under the Marie Skłodowska-Curie grant agreement No 801133. A portion of the manuscript was completed while S.M. was in residence as Ida Pfeiffer Guest Professor at the Department for Lithospheric Research, University of Vienna. M.B. acknowledges support provided by grants from the Carlsberg Foundation (CF18_1105) and the European Research Council (ERC Advanced Grant Agreement 833275 — DEEPTIME).

### Author contributions

I.G. conceived the idea and supervised the project, E.K. and I.G. designed the research, E.K. performed the confocal microscopy visualization, RNA encapsulation and DNA strand displacement experiments and associated fluorescence intensity analyses, I.P. prepared the initial setup, H.F. selected and provided the rock and mineral surfaces, H.F., E.K. and I.G. performed the SEM-EDX analyses of the surfaces, S.M. selected and provided the glasses and the associated whole rock analyses, M.B. provided the NWA 7533 specimen and prepared the sample to be suitable for the microscopy experiments. All authors contributed to the writing of the manuscript.

### Competing interests

The authors declare no competing financial interests.
